# A human papillomavirus 16 E2-TopBP1 dependent SIRT1-p300 acetylation switch regulates mitotic viral and human protein levels

**DOI:** 10.1101/2024.01.15.575713

**Authors:** Apurva T. Prabhakar, Claire D. James, Aya H. Youssef, Reafa A. Hossain, Ronald D. Hill, Molly L. Bristol, Xu Wang, Aanchal Dubey, Iain M. Morgan

## Abstract

An interaction between human papillomavirus 16 (HPV16) E2 and the cellular proteins TopBP1 and BRD4 is required for E2 plasmid segregation function. The E2-TopBP1 interaction promotes increased mitotic E2 protein levels in U2OS and N/Tert-1 cells, as well as in human foreskin keratinocytes immortalized by HPV16 (HFK+HPV16). SIRT1 deacetylation reduces E2 protein stability and here we demonstrate that increased E2 acetylation occurs during mitosis in a TopBP1 interacting dependent manner, promoting E2 mitotic stabilization. p300 mediates E2 acetylation and acetylation is increased due to E2 switching off SIRT1 function during mitosis in a TopBP1 interacting dependent manner, confirmed by increased p53 stability and acetylation on lysine 382, a known target for SIRT1 deacetylation. SIRT1 can complex with E2 in growing cells but is unable to do so during mitosis due to the E2-TopBP1 interaction; SIRT1 is also unable to complex with p53 in mitotic E2 wild type cells but can complex with p53 outside of mitosis. E2 lysines 111 and 112 are highly conserved residues across all E2 proteins and we demonstrate that K111 hyper-acetylation occurs during mitosis, promoting E2 interaction with Topoisomerase 1 (Top1). We also demonstrate that K112 ubiquitination promotes E2 proteasomal degradation during mitosis. The results present a model in which the E2-TopBP1 complex inactivates SIRT1 during mitosis and E2 acetylation on K111 by p300 increases, promoting interaction with Top1 that protects K112 from ubiquitination and therefore E2 proteasomal degradation.

**Importance:** Human papillomaviruses are causative agents in around 5% of all human cancers. While there are prophylactic vaccines that will significantly alleviate HPV disease burden on future generations, there are currently no anti-viral strategies available for the treatment of HPV cancers. To generate such reagents, we must understand more about the HPV life cycle, and in particular about viral-host interactions. Here we describe a novel mitotic complex generated by the HPV16 E2 protein interacting with the host protein TopBP1 that controls the function of the deacetylase SIRT1. The E2-TopBP1 interaction disrupts SIRT1 function during mitosis in order to enhance acetylation and stability of viral and host proteins. This novel complex is essential for the HPV16 life cycle and represents a novel anti-viral therapeutic target.

## Introduction

Human papillomaviruses (HPV) are causative agents in around 5% of all cancers and their life cycle depends upon epithelial differentiation (1, 2). The life cycle is tightly regulated via an intricate interaction between viral and host factors, and cancer results when this regulation is disrupted, preventing differentiation and promoting continued proliferation which results in accumulation of host cell DNA damage (3). The viral oncogenes E6 and E7 both contribute to the regulation of cell differentiation and proliferation via interaction with host factors, including p53 and pRb respectively (4). The viral genome is an 8kbp DNA episome whose replication is dependent upon the viral factors E1 and E2 interacting with host factors (5, 6).

E2 has a carboxyl terminus dimerization and DNA binding domain, binding to 12bp palindromic sequences in the viral genome that surround the A-T rich viral origin of replication (5). Following binding, the E2 amino terminal domain recruits the viral helicase E1 to the origin via a protein-protein interaction, whereupon E1 forms a di-hexameric helicase that activates viral replication in association with host polymerases (7, 8). In addition to viral replication, the E2 protein can also regulate viral and host gene transcription via interaction with host proteins, including BRD4 (9–12). A final role for E2 in the viral life cycle is segregation of the viral genome, where E2 acts as a “bridge” between the viral genome and host chromatin during mitosis to insure the viral genome resides in daughter nuclei following cell division (13). Initial studies with BPV1 E2 demonstrated that the host protein BRD4 was required for E2 interaction with mitotic chromatin and some studies indicated that this was also the mitotic chromatin receptor for HPV16 E2, while others suggested an alternative host factor was responsible for HPV16 E2 interaction with host mitotic chromatin (14–19).

We demonstrated that an interaction between HPV16 E2 (from now on E2 will mean HPV16 E2 unless stated otherwise) and TopBP1 is essential for E2 interaction with mitotic chromatin and E2 plasmid segregation function, and that the E2-TopBP1 interaction is dependent upon E2 phosphorylation on serine 23 by CK2 (20–22). More recently we demonstrated that an E2 interaction with both TopBP1 and BRD4 is required for E2 interaction with mitotic chromatin and E2 plasmid segregation function (23). During these studies we observed a stabilization of E2 during mitosis that is dependent upon an interaction with TopBP1 but not BRD4. In addition, TopBP1 was also stabilized in an E2 interaction dependent manner. These observations were made in the hTERT immortalized human foreskin keratinocytes N/Tert-1 and U2OS cells, as well as in human foreskin keratinocytes immortalized by HPV16.

Previously, we demonstrated that the class III deacetylase SIRT1 can deacetylate E2, and that this deacetylation reduces E2 protein stability, while others have demonstrated a critical role for SIRT1 during the HPV31 life cycle (24, 25). SIRT1 can also regulate the acetylation status and function of TopBP1 (26, 27). Here we demonstrate that, during mitosis in N/Tert-1 and U2OS cells, the interaction between E2 and SIRT1 is disrupted in an E2-TopBP1 interacting dependent manner and that E2 acetylation is increased. E2 acetylation occurs on lysine 111 (K111) and this acetylation promotes interaction with Topoisomerase 1 (Top1) and protects lysine 112 (K112) from ubiquitination. This explains why acetylation of K111 promotes E2 stabilization during mitosis. We demonstrate aberrant interaction of the E2 K mutants with mitotic chromatin, and that the K111R mutant is defective in transient replication and transcription assays. Introduction of the E2 K mutants into the genome abrogated the immortalization potential of HPV16. SIRT1 is also inactive, in an E2-TopBP1 interacting dependent manner, on p53 during mitosis resulting in enhanced p53 expression and acetylation. In HFK+HPV16 cells, SIRT1 expression is reduced at the protein level during mitosis and there is increased mitotic expression of E2 and p53 in these cells; a similar phenotype to that in the N/Tert-1 and U2OS cells expressing E2 only. Significantly, isogenic HFK immortalized by only E6 and E7 (HFK+E6E7) do not show this phenotype; SIRT1 is unaffected and there is no increased acetylation and expression of p53 in these cells during mitosis. We also demonstrate that p300 is the acetylase responsible for the increased acetylation of viral and host proteins during mitosis in the HFK+HPV16 cells. The results demonstrate that an E2-TopBP1 interaction controls SIRT1 activity during mitosis and that this increases p300 acetylation of viral and cellular proteins important during the HPV16 life cycle.

## Results

### An E2-TopBP1 interaction turns off SIRT1 function during mitosis

Previously, we demonstrated increased expression of E2 and TopBP1 during mitosis in HFK+HPV16 cells, and that SIRT1 deacetylation of E2 reduces E2 protein expression (21, 24). We therefore investigated whether the increased expression of E2 and TopBP1 during mitosis in HFK+HPV16 was due to increased E2 acetylation via disruption of SIRT1 mitotic function. Figure 1A demonstrates that SIRT1 protein expression is reduced in mitotic HFK+HPV16 cells (lane 4) but not in isogenic HFK immortalized by E6E7 (lane 6), nor in N/Tert-1 cells that have been G418 selected (Vec, for Vector control, lane 2). The previously reported increase in E2 and TopBP1 was also detected in the HFK+HPV16 positive cells (lane 4). Mitotic cell enrichment was carried out by double thymidine blocking (DTB) followed by a release for 19 hours (HFK) and 16 hours (N/Tert-1 Vec), and mitotic enrichment is demonstrated by increased cyclin B expression (compare lanes 2, 4 and 6 with 1, 3 and 5). In mitotic N/Tert-1 and HFK+E6E7 cells there is an increase in SIRT1 during mitosis (lanes 2 and 6) in sharp contrast to the reduction observed in HFK+HPV16 cells (lane 4), and no increase in TopBP1 expression. An acetyl lysine co-immunoprecipitation determined that the acetylation of E2, TopBP1 and p53 was increased during mitosis (Figure 1B). Lane 1 is a co-IP with the control HA antibody, and it does not interact with any of the proteins under study. In N/Tert-1 and HFK+HPV16 cells p53 is acetylated at time 0h (0 hours) (lanes 2 and 4) and this acetylation is increased in mitotic HFK+HPV16 cells (lane 5) and not in N/Tert-1 cells (lane 3) where p53 expression is reduced (Figure 1A). p53 levels in HFK+HPV16 cells are detectable as E6 is spliced to E6* which removes the p53 degradation domain of E6 (28). We have demonstrated abundant p53 expression in HPV positive head and neck cancer cell lines and patient derived xenografts, as well as in primary keratinocytes immortalized by the entire HPV16 genome (29). No p53 is detectable in the HFK+E6E7 cells as E6 is not spliced and can therefore target p53 for proteasomal degradation. The acetylation of E2 is detectable in 0h HFK+HPV16 cells and this increases during mitosis (lanes 4 and 5), reflective of the increased E2 protein expression (Figure 1A). There is no detectable acetylation of the residual levels of SIRT1 in mitotic HFK+HPV16 (lane 5). Next we investigated whether the residual SIRT1 in mitotic cells was able to interact with TopBP1, E2 and p53 using a SIRT1 co-immunoprecipitation (Figure 1C). TopBP1 interaction with SIRT1 is not disrupted in mitosis in any of the samples. In HFK+HPV16 cells, SIRT1 interaction with E2 and p53 is disrupted during mitosis (lane 5). Figure S1 quantitates repeats of Figure 1A demonstrating significant changes in TopBP1, SIRT1, E2 and p53 between the samples.

**Figure 1.**
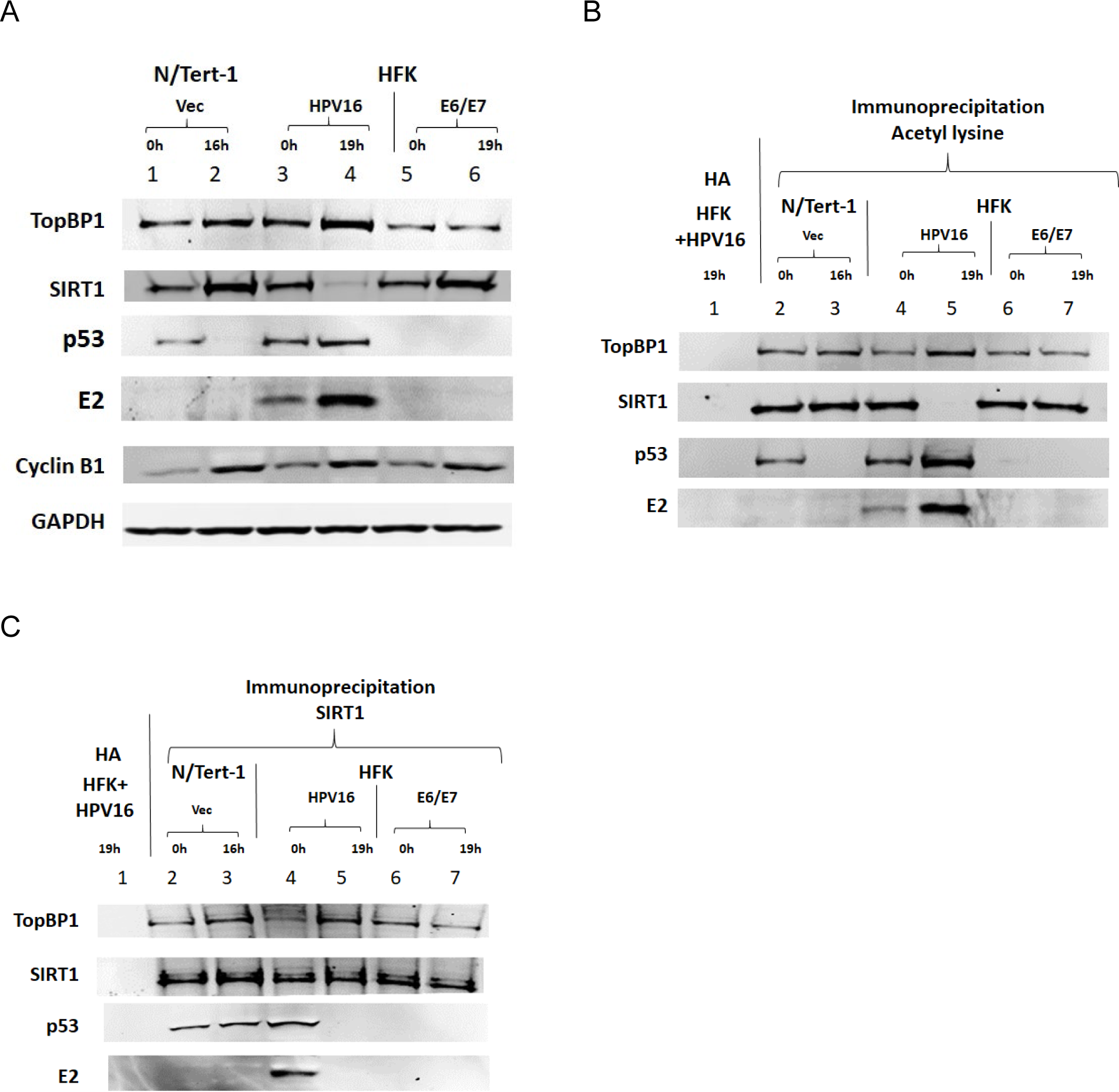
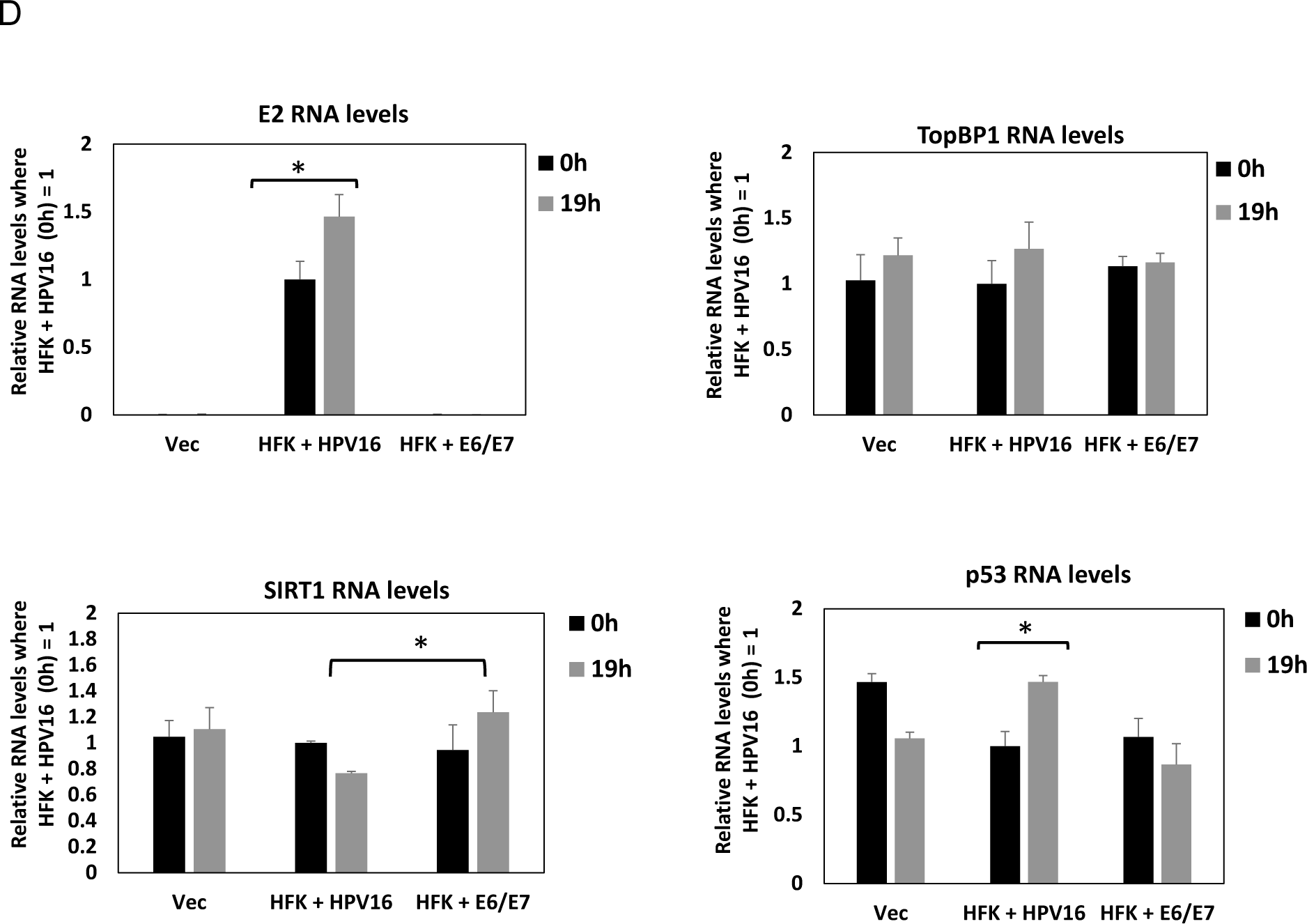
Increased expression of E2, TopBP1 and p53 in mitotic enriched HFK+HPV16 cells. A. N/Tert-1 (lanes 1 and 2), HFK+HPV16 (lanes 3 and 4) and HFK+E6/E7 (lanes 5 and 6) were arrested by double thymidine block (DTB) (lanes 1, 3, 5) and released for 16 hours (N/Tert1, lane 2) or 19 hours (HFK+HPV16 and HFK+E6/E7, lanes 4 and 6, respectively) to enrich for mitotic cells (confirmed by enhanced expression of cyclin B1). Western blotting demonstrated expression of the indicated proteins. B and C. The extracts in A were immunoprecipitated with an acetyl-lysine antibody (lanes 2-7) or an HA control antibody with HFK+HPV16 extract (lane 1). Western blotting demonstrated the level of acetyl-lysine pull-down of the indicated proteins. C. The extracts in A were immunoprecipitated with a SIRT1 antibody (lanes 2-7) or an HA control antibody with HFK+HPV16 extract (lane 1). Western blotting demonstrated the level of SIRT1 pull-down of the indicated proteins. D. RNA levels of the indicated genes were determined in DTB arrested and mitotic enriched cells. Significant changes are indicated with *, p-value < 0.05.

One of the functions of SIRT1 during mitosis is to deacetylate histones and promote chromatin condensation (30, 31). The reduction in SIRT1 function during mitosis could therefore alter transcription from the viral and host genome in HFK+HPV16 cells. Figure 1D demonstrates that there is a significant increase in E2 and p53 RNA levels during mitosis in HFK+HPV16 cells; p53 RNA levels are not increased in mitosis in either N/Tert-1 Vec or HFK+E6E7 cells. With SIRT1 there is a reduction in mitotic HFK+HPV16 cells and an increase in HFK+E6E7, although neither reached significance. When paired, there is a significant difference between mitotic SIRT1 RNA levels in HFK+HPV16 cells versus HFK+E6E7 cells. There is no significant change in the SIRT1 levels in N/Tert-1 Vec cells. Compared to the levels of protein changes observed (Figure 1A), the changes in RNA levels were minimal although may contribute to the changes in protein levels observed in mitosis (Figure 1A and Figure S1). There is no significant change in TopBP1 RNA levels in any of the samples.

As the changes in p53 and SIRT1 levels were not observed in HFK+E6E7 (Figure 1), and E2 increases TopBP1 protein levels during mitosis (21, 22), the ability of E2 to regulate SIRT1 function during mitosis via interaction with TopBP1 was investigated in N/Tert-1 and U2OS cells (Figure 2). Figure 2A demonstrates that in N/Tert-1 cells stably expressing E2-WT there is an increase in TopBP1, E2 and p53 during mitosis, which is similar to the HFK+HPV16 cells (Figure 1A). An E2 mutant that cannot bind TopBP1 (E2-S23A), did not increase the expression of TopBP1, p53 or E2 during mitosis (compare lane 6 with lane 4). An acetyl lysine immunoprecipitation demonstrated increased acetylation of TopBP1 and E2 during mitosis, and a slight increase in acetylated p53 (compare lane 5 with lanes 3 and 7). In both N/Tert-1 Vec and E2-S23A cells, there is a reduction in p53 protein levels during mitosis (Figure 2A). Unlike HFK+HPV16 cells, E2-WT did not reduce SIRT1 protein levels in mitosis when compared with non-mitotic cells (Figure 2A, lanes 3 and 4), although SIRT1 protein levels did not increase as they do in Vec and E2-S23A cells (lanes 2 and 6, Figure 2A). However, as in HFK+HPV16 cells there is a reduction in SIRT1 acetylation levels during mitosis only in the presence of E2-WT (Figure 2B, lane 5). Figure 2C demonstrates that the ability of SIRT1 to complex with p53 and E2 during mitosis is abrogated only in E2-WT cells, as it is in HFK+HPV16 cells. Figure S2 quantitates repeats of Figure 2A demonstrating significant changes in TopBP1, SIRT1, E2 and p53 between the samples.

**Figure 2.**
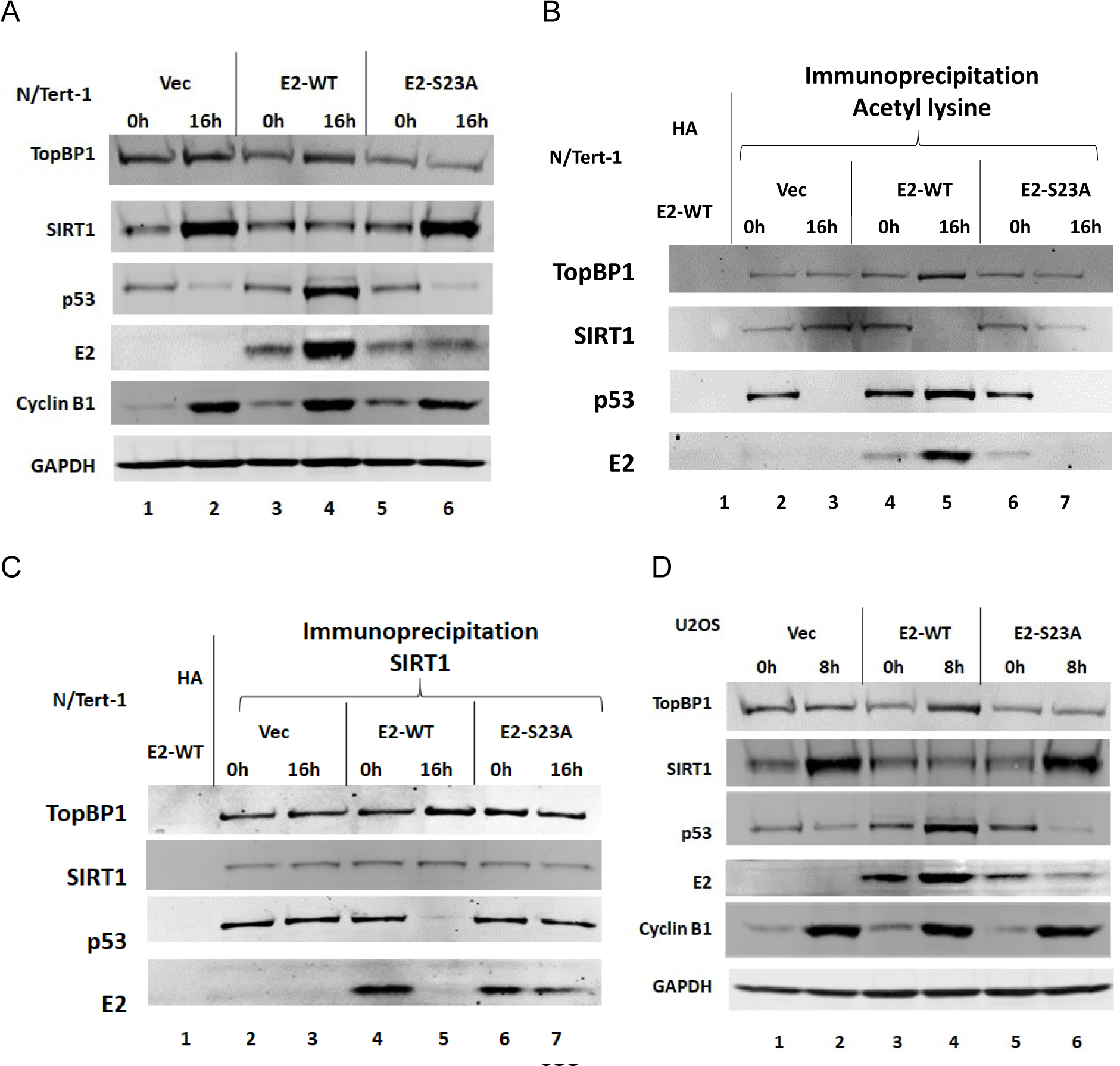
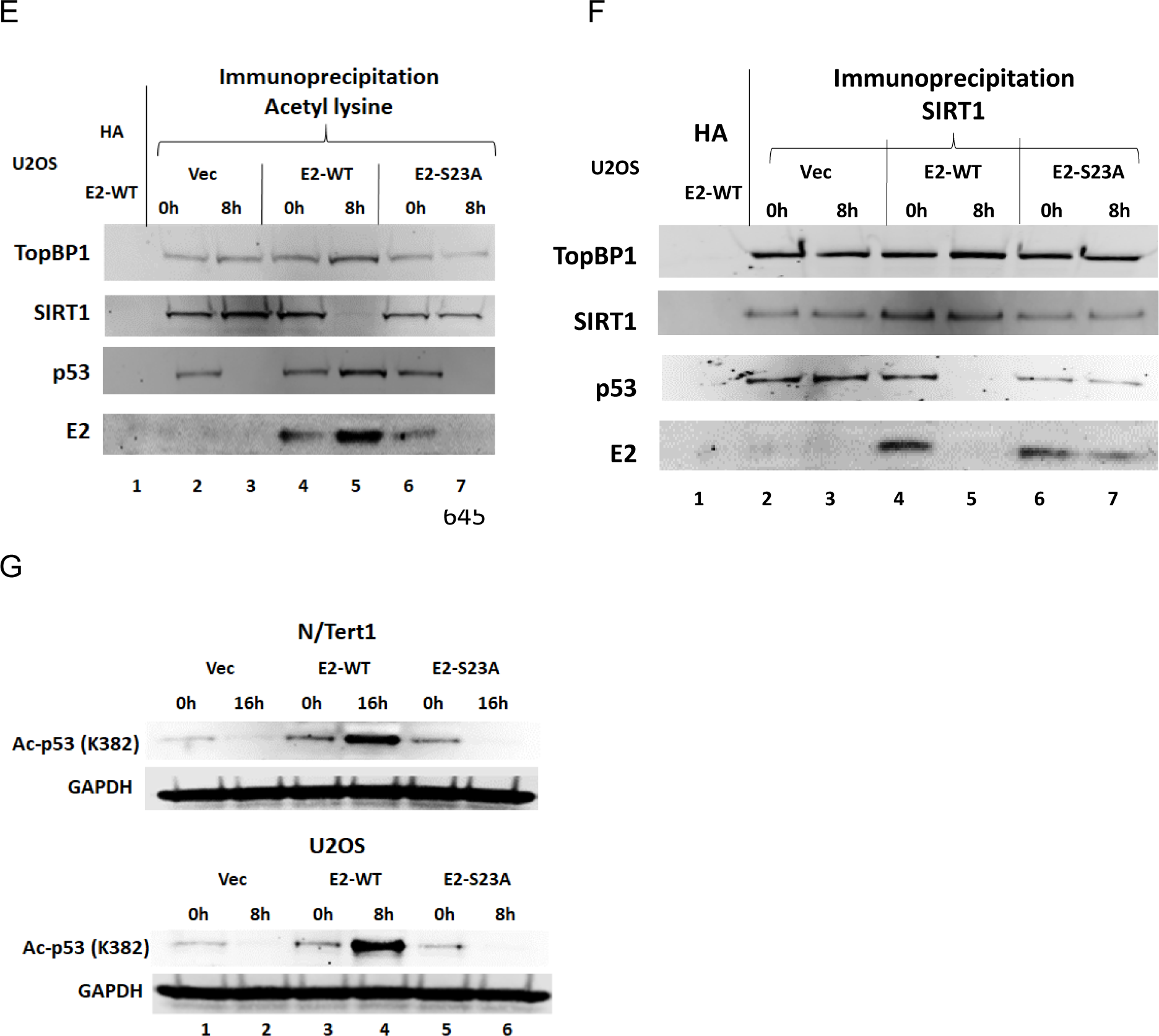
An E2-TopBP1 interaction is required to turn-off SIRT1 function and increase acetylation during mitosis. A. N/Tert-1 Vec (pcDNA vector control) (lanes 1 and 2), N/Tert-1 E2-WT (lanes 3 and 4) and N/Tert-1 E2-S23A (TopBP1 binding interacting mutant, lanes 5 and 6) were DTB (lanes 1, 3 and 5) and released for 16 hours to enrich for mitotic cells (lanes 2, 4 and 6). Western blotting demonstrated expression of the indicated proteins. B. The extracts in A were immunoprecipitated with an acetyl-lysine antibody (lanes 2-6) or an HA control antibody with E2-WT extracts (lane 1). Western blotting demonstrated the level of acetyl-lysine pull-down of the indicated proteins. C. The extracts in A were immunoprecipitated with a SIRT1 antibody (lanes 2-7) or an HA control antibody with HFK+HPV16 extract (lane 1). Western blotting demonstrated the level of SIRT1 pull-down of the indicated proteins. D-F represent a repeat of A-C in U2OS cells. G. The extracts in A and D were western blotted with an antibody specific for p53 acetylation on lysine 382 (AcK382), a residue known to be deacetylated by SIRT1.

Figures 2D to 2F established that, in U2OS cells, E2-WT has the same phenotype as in N/Tert-1 cells (repeats of Figure 2D are quantitated in Figure S3). E2, TopBP1, SIRT1 and p53 RNA levels were not significantly different during mitosis in the Vec, E2-WT and E2-S23A N/Tert-1 (Figure S4) or U2OS (Figure S5) cells when compared with non-mitotic cells. To confirm that SIRT1 is not deacetylating target lysines on p53, western blots with a p53 K382 acetylated antibody (Ac-p53 (K382)) confirmed that the SIRT1 target residue had increased acetylation during mitosis in N/Tert-1 and U2OS cells, confirming the “knock-out” of SIRT1 function by E2-WT (Figure 2G). The p53 K382 acetylation is quantitated in Figure S2 (N/Tert-1 cells) and Figure S3 (U2OS cells).

The results in Figure 2 demonstrate that E2 manipulates SIRT1 function during mitosis in a TopBP1 interaction dependent manner, and Figure 1 confirms this happens in HFK+HPV16 cells. E2 interaction with TopBP1 is required for increased p53 acetylation on lysine 382, an acetylation known to stabilize p53. We next determined the E2 residues that regulate E2 acetylation and stability during mitosis.

### E2 K111 and K112 regulate the stability of E2 during mitosis

E2 lysines 111 and 112 (K111 and K112) are highly conserved across HPV types, and HPV16 E2 K111 is acetylated (32). A panel of N/Tert-1 cells were established stably expressing K111 and K112 E2 mutants that mimicked acetylation (K to Q) or mimicked non-acetylation (K to R) (Figure 3A). Using DTB and release for 16 hours, mitotic enrichment was carried out and western blotting demonstrated the expression of the E2 proteins and cyclin B expression demonstrated mitotic enrichment (Figure 3B). E2-WT had increased expression during mitosis (lane 4 and 12), E2-K111R (mimicking non-acetylation) had no detectable expression during mitosis (lane 6), while E2-K111Q (mimicking acetylation) expressed similarly to E2-WT during mitosis (compare lane 14 with lane 12). E2-K112R levels increased during mitosis similarly to E2-WT (compare lane 8 with lane 4). E2-K111R+K112R (double lysine mutant) protein levels did not increase during mitosis but were detectable. All of the mutants were able to complex with TopBP1, if they were detectable in the input blot (Figure 3C). An acetyl-lysine immunoprecipitation demonstrated increased acetylation of E2-WT and E2-K112R during mitosis, none of the other mutants were pulled down by the acetyl-lysine antibody, including K111R in non-mitotic cells (lane 6) (Figure 3D). The results demonstrate that there is one major site of acetylation on E2 in and out of mitosis, K111. Acetylation of K111 promotes interaction with Topoisomerase 1 (Top1) (32, 33) and we investigated the ability of the E2 mutants to interact with Top1 (Figure 3E). E2-K111R and E2-K111R+K112R were unable to complex with Top1 (lanes 6 and 7, and 17 and 18, respectively). The significant mitotic increase of E2-WT, K112R, K111Q protein expression, as well as TopBP1 in these three samples, when compared with the time 0h sample (Figure 3B) was confirmed using multiple sample blots (Figure S6). E2 nor TopBP1 RNA levels were changed in mitosis in any of the cell lines, nor were there any changes in expression due to the E2 mutations between samples (Figure S7).

**Figure 3.**
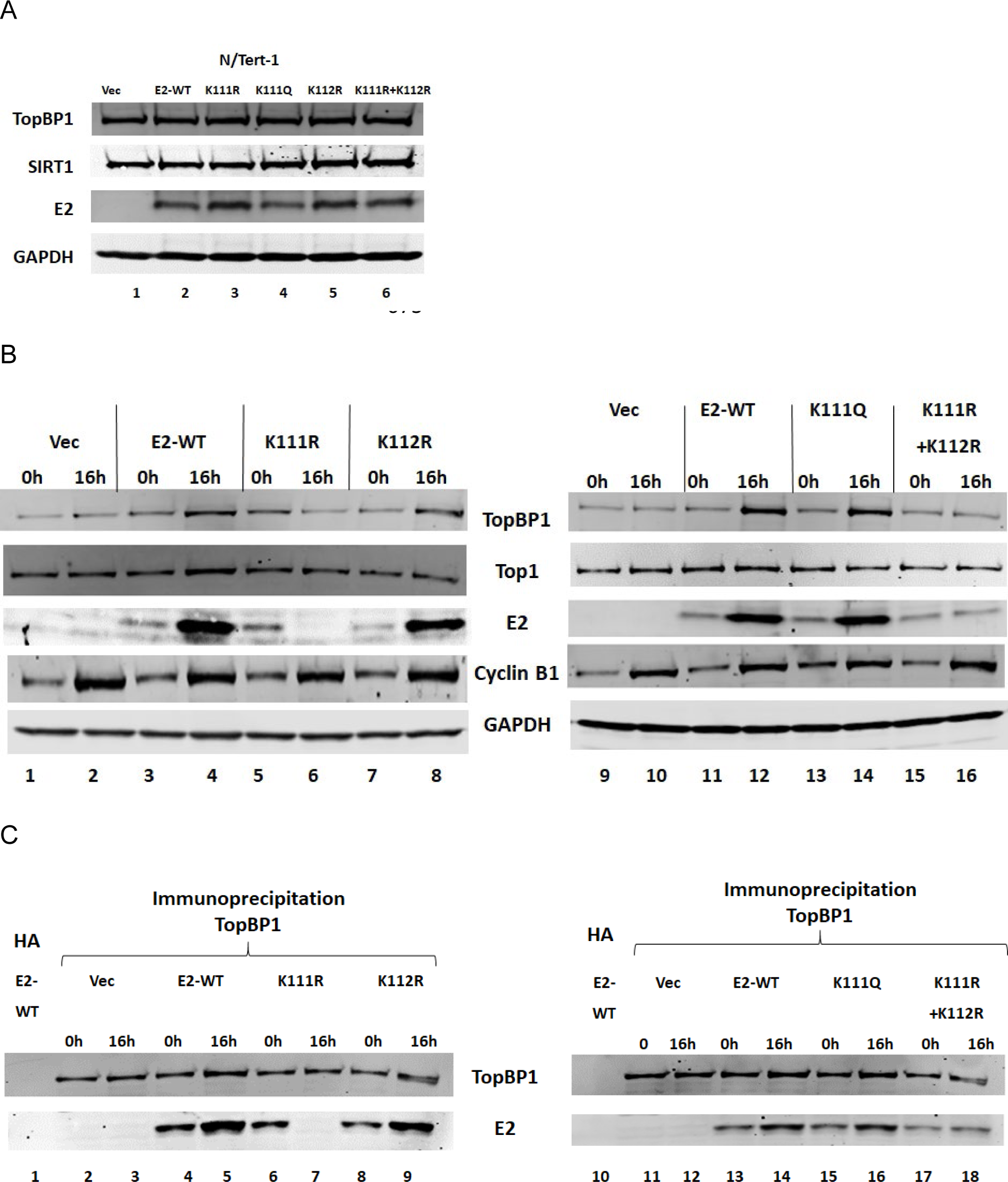
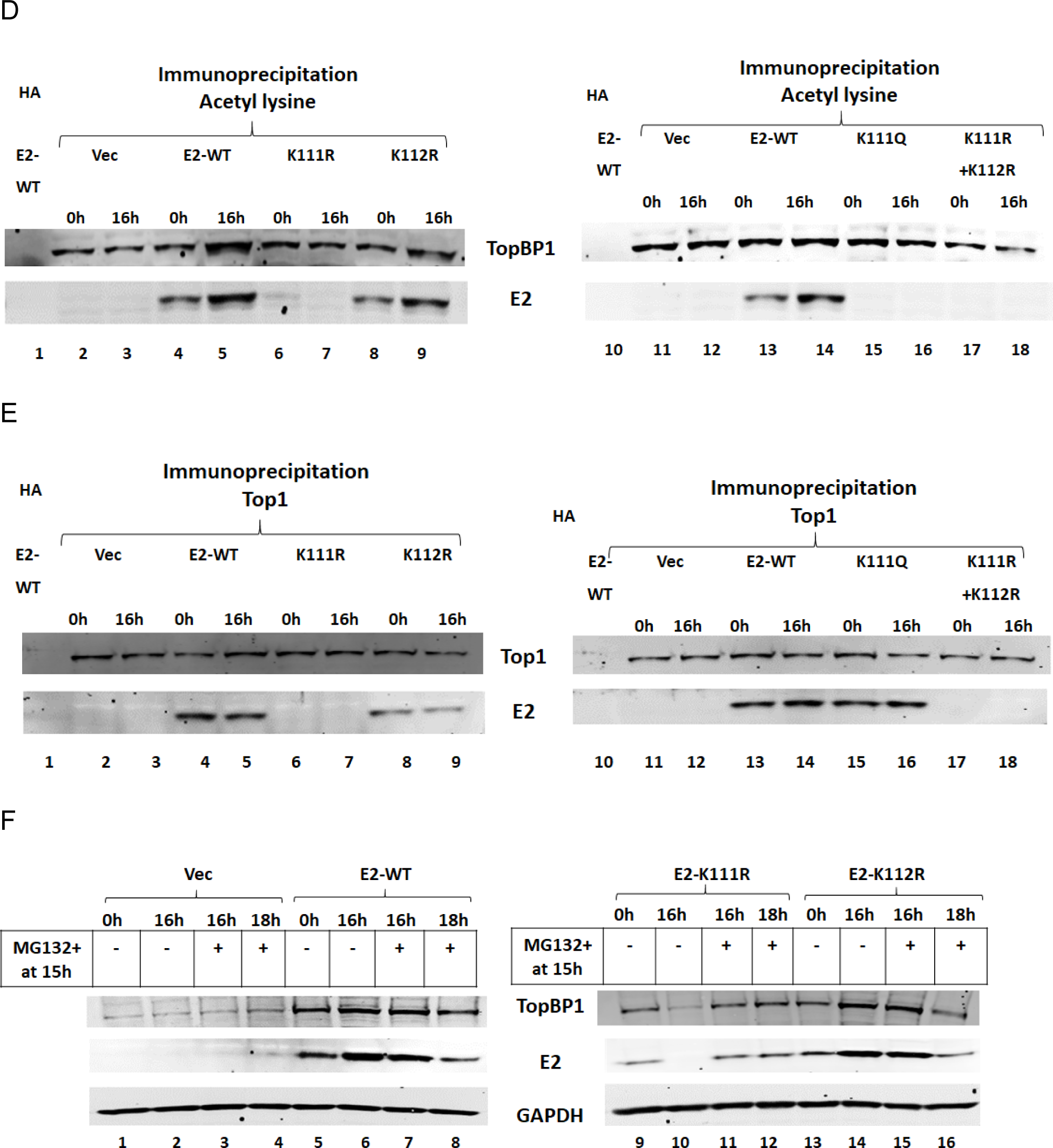
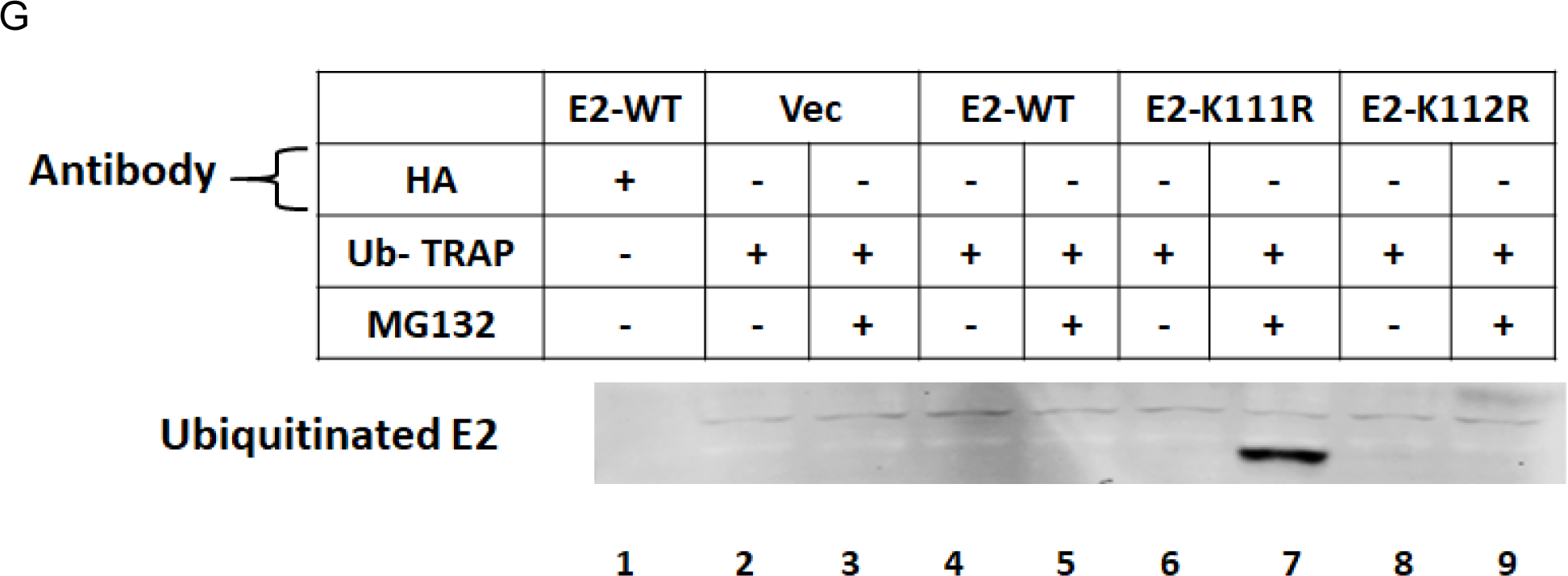
E2 lysine 111 (K111) and 112 (K112) regulate E2 stability during mitosis. A. N/Tert-1 cells stably expressing the E2 proteins indicated were generated. B. Cell lines were DTB treated (lanes 1, 3, 5, 7, 9, 11, 13, 15) or DTB and released for 16 hours to enrich for mitotic cells (lanes 2, 4, 6, 8, 10, 12, 14, 16). Western blotting demonstrated expression of the indicated proteins. C. The extracts in B were immunoprecipitated with a TopBP1 antibody (lanes 2-9 and 11-18) or an HA control antibody with E2-WT extracts (lanes 1 and 10). Western blotting demonstrated the level of TopBP1 pull-down of the indicated proteins. D. The extracts in B were immunoprecipitated with an acetyl-lysine antibody (lanes 2-9 and 11-18) or an HA control antibody with E2-WT extracts (lanes 1 and 10). Western blotting demonstrated the level of acetyl-lysine pull-down of the indicated proteins. E. The extracts in B were immunoprecipitated with a Top1 antibody (lanes 2-9 and 11-18) or an HA control antibody with E2-WT extracts (lanes 1 and 10). Western blotting demonstrated the level of Top1 pull-down of the indicated proteins. F. The indicated cell lines were DTB treated (lanes 1, 5, 9 and 13) or released for 16 hours to enrich for mitotic cells (lanes 2, 6, 10 and 14). MG132 was added at 15 hours (lanes 3, 4, 7, 8, 11, 12, 15, 16) following DTB released and harvested either one hour (lanes 3, 7, 11, 15) or 3 hours (lanes 4, 8, 12, 16) later. Western blotting demonstrated the level of the indicated proteins. G. The 16 hour release samples from F + or – MG132 were tested for E2 ability to be ubiquitinated using a ubiquitin trap (see materials and methods).

The recovery of E2K111R+K112R protein expression in mitosis, when compared with E2-K111R, suggested that K112 was ubiquitinated, targeting the E2 protein for proteasomal degradation. To test this, the proteasomal inhibitor MG132 was added 15 hours following release from the DTB and E2 protein levels determined at 16 and 18 hours following DTB release (Figure 3F). Without addition of MG132, E2-K111R was undetectable during mitosis, similarly to Figure 3B. However, addition of MG132 for 1 hour prior to cell harvesting resulted in detection of E2-K111R in mitotic cells, demonstrating that proteasomal inhibition restored expression (lane 11). Interestingly, E2-WT levels 3 hours following the addition of MG132 are reduced, suggesting the decrease of E2 protein levels following exit from mitosis may not be mediated by proteasomal degradation. To determine whether E2-K111R has increased ubiquitination in mitotic cells we used the Chromo Tek Ubiquitin-Trap (Figure 3G). The samples assayed were mitotic samples treated with and without MG132 for 1h prior to protein extraction (Figure 3F). Lane 7 demonstrates that in mitotic cells treated with MG132 E2-K111R is ubiquitinated, which would target the protein for proteasomal turnover, explaining the lack of E2-K111R protein expression during mitosis. The results from Figure 1-3 support a model in which SIRT1 function is, at least partially, inactivated during mitosis in an E2-TopBP1 interacting dependent manner. This SIRT1 inactivation results in enhanced acetylation of E2 K111, promoting interaction with Top1. This interaction protects E2 K112 from ubiquitination and proteasomal turnover during mitosis promoting increased E2 protein levels.

### Lysine mutations in E2 disrupt E2 mitotic interaction and abolish HPV16 primary cell immortalization

To determine the cellular localization of the E2 lysine mutants, stable U2OS cells expressing the mutants were generated (Figure 4A). Increased E2 expression and acetylation occurs in U2OS cells in an identical manner to that in N/Tert-1 cells (Figure 2), and imaging mitotic U2OS cells is more straightforward than imaging mitotic N/Tert-1 cells. Figure 4B represents examples of mitotic U2OS cells stably expressing the indicated mutants. In the absence of E2, TopBP1 is located on mitotic chromatin (Vec, top panels) and WT-E2 is co-located with TopBP1 on mitotic chromatin (second panel down), as previously reported (21). E2-K111R expression during mitosis is not detected by western blotting and in immunofluorescence there was a reduced E2-K111R signal in mitotic cells (third panel down). The interaction of TopBP1 with mitotic chromatin is not affected by E2-K111R expression. E2-K111Q is located on mitotic chromatin (fourth panel down), but this mutant removes TopBP1 from mitotic chromatin. E2 K112R, well expressed during mitosis, fails to locate to mitotic chromatin (fifth panel down) and does not disrupt the ability of TopBP1 mitotic interaction. E2-K111R+K112R has a similar phenotype as E2-K112R (sixth panel down). In interphase cells, E2-K111R was detected in the cytoplasm when compared with E2-WT, and E2-K112R and E2-K111R+K112R also had more cytoplasmic E2 staining. The levels of E2 and TopBP1 interaction with mitotic chromatin were determined using the Keyence imaging system over multiple mitotic cells and confirm the phenotypes observed in Figure 4B (Figure 4C). Nuclear and cytoplasmic E2 staining was also quantitated and confirmed significantly more cytoplasmic staining for E2-K111R, E2-K112R and E2-K111R+K112R (Figure 4D).

**Figure 4.**
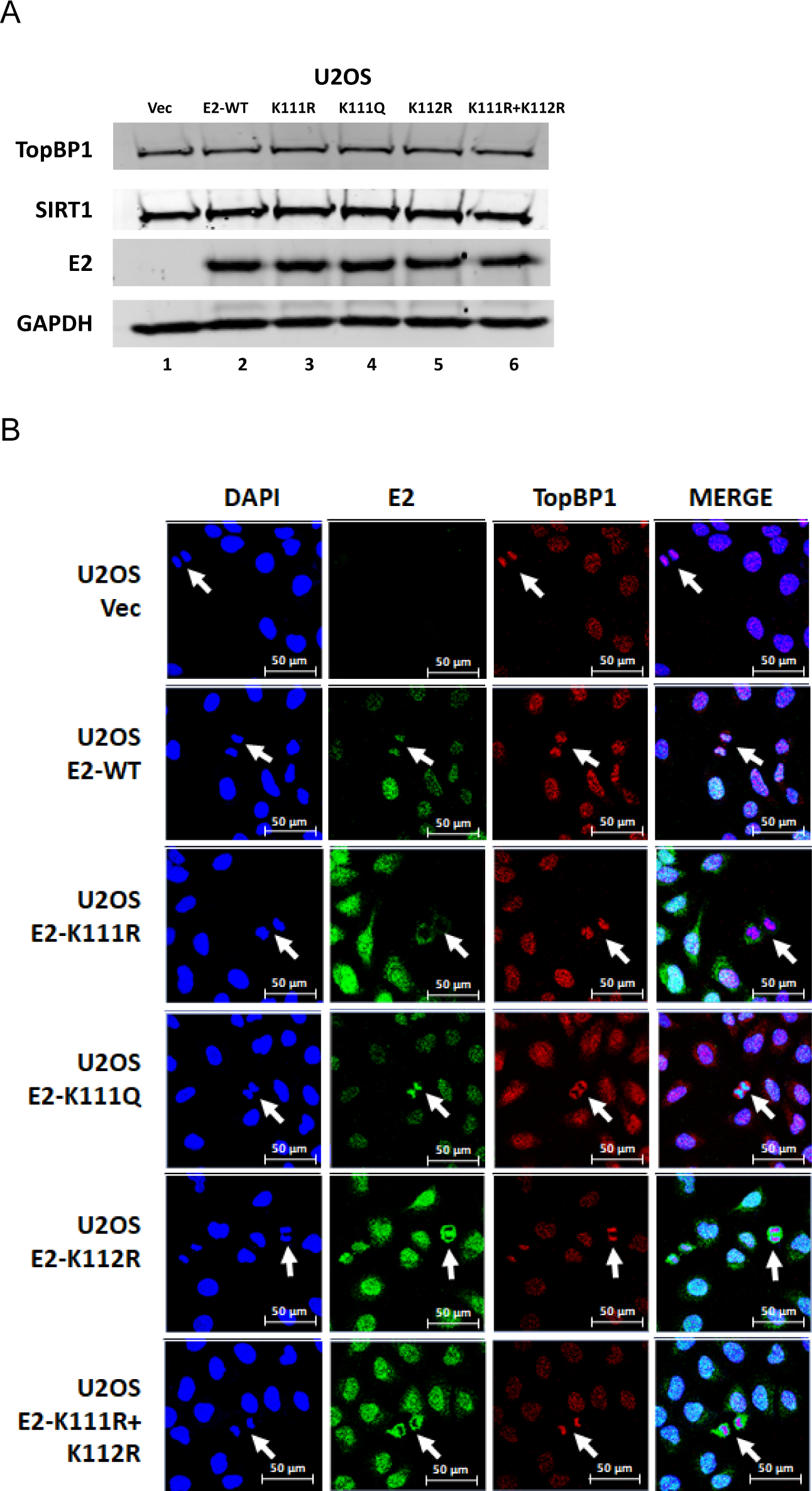
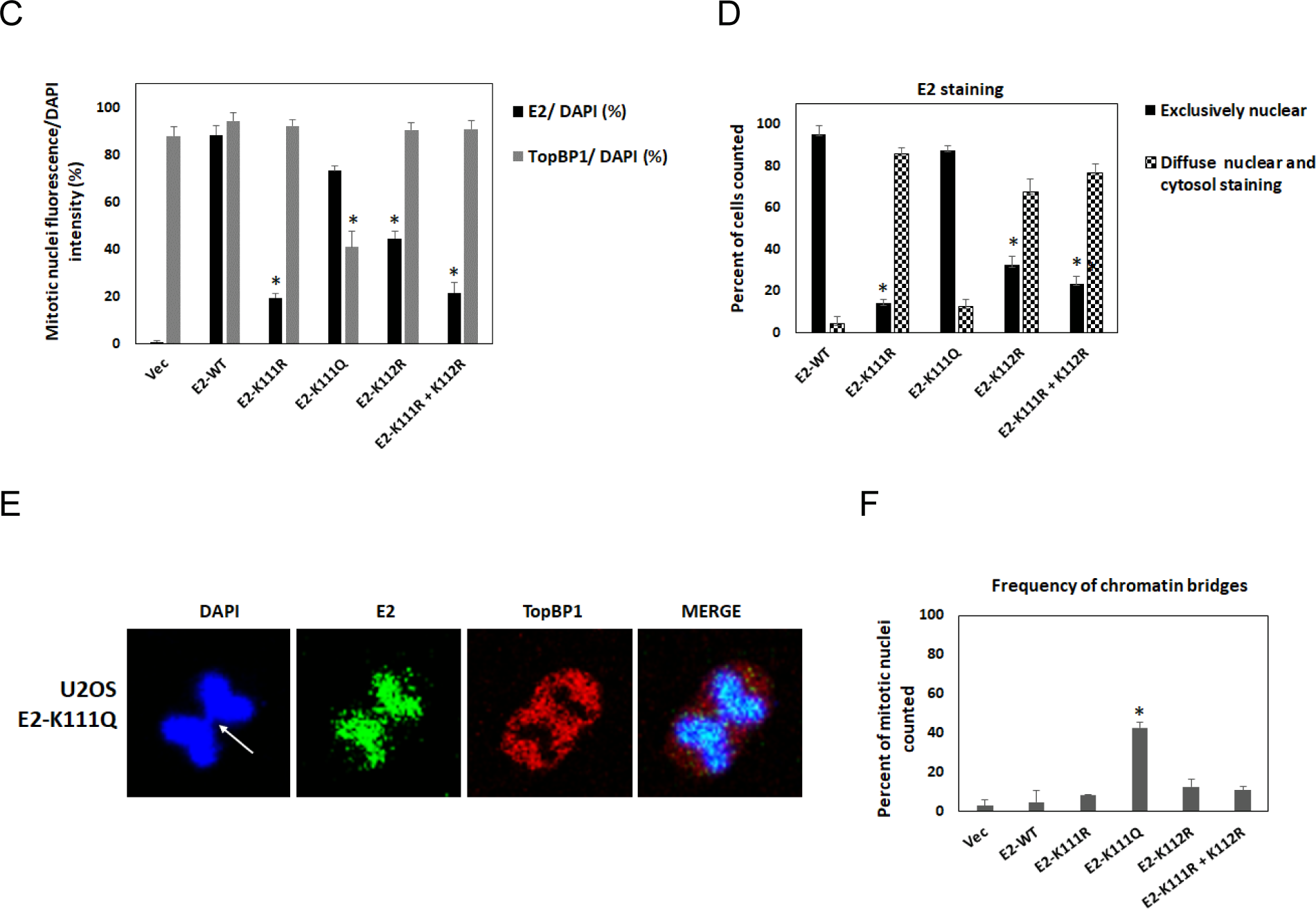
Aberrant interaction of E2 lysine mutants with mitotic chromatin. A. U2OS cells stably expressing the indicated E2 proteins were generated. Western blotting demonstrated expression of the indicated proteins. B. Cells were grown and fixed without enrichment for mitotic cells. Random mitotic cells are indicated with white arrows and E2 (green) and TopBP1 (red) staining carried out. C. The number of mitotic bodies that retained E2 or TopBP1 expression were determined using a Keyence imaging system. D. The Keyence system also determined the nuclear/cytosolic localization of the indicated E2 proteins. E. The E2 K111Q mutant complexed with mitotic chromatin but removed TopBP1 generating chromatin bridges (quantitated in F).

TopBP1 is an active protein during mitosis and is critical for maintaining genome integrity (16, 34–43). The ability of E2-K111Q to remove TopBP1 from mitotic chromatin indicated this protein may disrupt mitosis. We investigated the number of mitotic cells that contained anaphase bridges and observed an increased number of cells with this phenotype in the E2-K111Q cells, when compared with control and other E2 expressing cells. Figure 4E provides an enhanced image of one of the cells containing an anaphase bridge, and Figure 4F provides a quantitation of the occurrence, demonstrating a significant increase in E2-K111Q cells.

The ability of the E2 K mutants to immortalize human foreskin keratinocytes (HFK) was investigated. HPV16 genomes were generated with E2 mutations encoding K111R, K111Q and K112R, and along with wild type HPV16 genomes were transfected into two independent primary foreskin keratinocyte donor cells. In both donors HPV16 wild type generated immortalized clones that were grown out into cell lines; none of the K mutants were able to generate clones or immortalized lines (Table 1). This demonstrates that wild type E2-K111 and K112 are critical for the immortalization of HFK. Androphy and colleagues demonstrated that K111 is critical for the transcription and replication function of E2 and we observed the same phenotype (Figure S8), all mutants except for K111R were similar to wild type in their transcription and replication function in C33a cells. The K111R phenotype could be related to a failure to complex with Top1 (Figure 3C and (32)).

**Table 1.** Mutation of E2 lysine 111 or 112 abrogates the immortalization potential of HPV16. Two independent human foreskin donors were transfected with the indicated HPV16 genomes and the subsequent growth of colonies and immortalized cell lines determined. K111R = E2 lysine 111 mutated to arginine; K111Q = E2 lysine 111 mutated to glutamine; K112R = E2 K112 mutated to arginine.

|  | HPV16<br>WT |  | HPV16<br>K111R |  | HPV16<br>K111Q |  | HPV16<br>K112R |  |
| --- | --- | --- | --- | --- | --- | --- | --- | --- |
|  | Colonies | Immortal? | Colonies | Immortal? | Colonies | Immortal? | Colonies | Immortal? |
| Donor 1 | **6** | **Yes** | 0 | No | 0 | No | 0 | No |
| Donor 2 | **3** | **Yes** | 0 | No | 0 | No | 0 | No |

### p300 is the predominant mitotic acetylase for E2

p300 is a histone acetyl transferase involved in the acetylation and functional regulation of E2 proteins (32, 33, 44–49). Knockdown of p300 in N/Tert-1 cells abolished the increase of E2 and TopBP1 mitotic protein levels (Figure 5A, compare lane 4 with lane 8). An acetyl lysine immunoprecipitation demonstrated that p300 is required for detectable E2 acetylation in and out of mitosis (Figure 5B, compare lanes 4 and 5 with 8 and 9). For TopBP1 there is a pulldown in all lanes demonstrating that p300 is not the only acetylase that can target TopBP1, although the increase in mitosis in the E2 expressing cells is mediated by p300 (compare lane 5 with 9). There is increased SIRT1 acetylation during mitosis in the E2-TopBP1 samples following p300 knockdown (Figure 5B, compare lanes 5 and 9). This demonstrates that the E2-TopBP1 stabilization due to p300 acetylation prevents SIRT1 acetylation by another acetylase, and that this control is lost following p300 knockdown due to the reduction in E2-TopBP1 protein levels. p300 knockdown does not alter SIRT1 acetylation levels in N/Tert-1 Vec control cells demonstrating that p300 is not a SIRT1 acetylase. p300 is also acetylated but not altered by E2 expression or cell cycle changes (lanes 2-5). These experiments were repeated with an additional p300 siRNA and generated essentially the same results (Figure S9).

**Figure 5.**
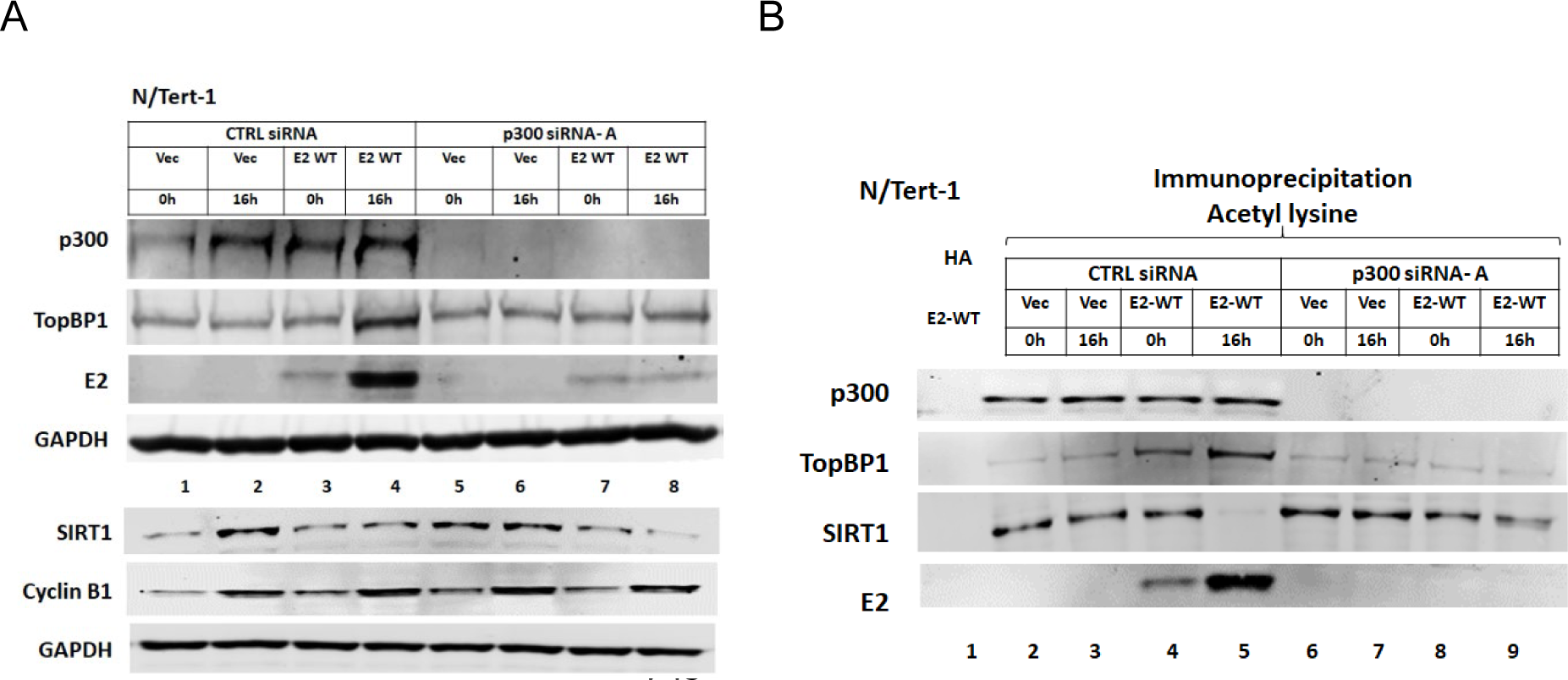
p300 enhances E2 and TopBP1 acetylation in mitosis in N/Tert-1 cells. A. N/Tert-1 Vec (lanes 1, 2, 5, 6) and N/Tert-1 E2-WT (lanes 3, 4, 7, 8) were DTB treated (lanes 1, 3, 5, 7) or released for 16 hours to enrich for mitotic cells (lanes 2, 4, 6, 8). Cells were treated with control siRNA (lanes 1-4) or siRNA targeting p300 (lanes 5-8). Western blotting demonstrated expression of the indicated proteins. B. The extracts in A were immunoprecipitated with an acetyl-lysine antibody (lanes 2-9) or an HA control antibody with E2-WT extracts (lane 1). Western blotting demonstrated the level of acetyl-lysine pull-down of the indicated proteins.

Next, we investigated whether p300 was responsible for E2 mitotic acetylation in HFK+HPV16 cells (Figure 6A). Following p300 knockdown the increase in E2, TopBP1 and p53 levels during mitosis is abolished (compare lane 4 with lane 8); p300 is a known p53 acetylase (50). An acetyl lysine IP (Figure 6B) demonstrated that acetylation of p53 and E2 was reduced following p300 knockdown, and that p300 is not the SIRT1 acetylase. We repeated these experiments with an additional p300 siRNA and got essentially identical results (Figure S10). Overall, the results mimic those observed in the N/Tert-1+E2 cells, including the regulation of SIRT1 acetylation (Figure 5).

**Figure 6.**
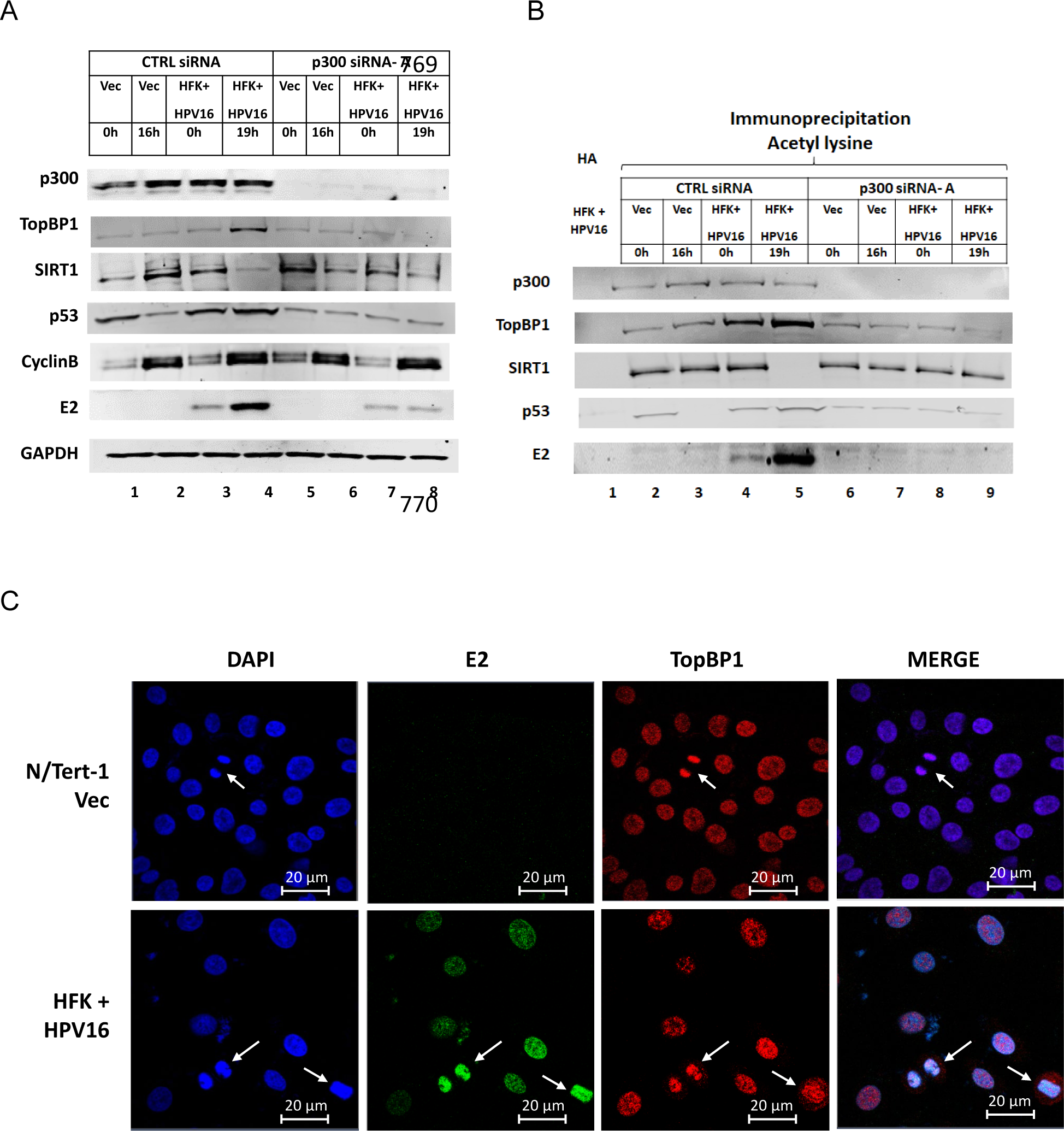
p300 enhances E2 and TopBP1 acetylation in mitosis in HFK+HPV16 cells. A. N/Tert-1 Vec (lanes 1, 2, 5, 6) and HFK+HPV16 (lanes 3, 4, 7, 8) were DTB treated (lanes 1, 3, 5, 7) or released for 16 hours (lanes 2 and 6) or 19 hours (lanes 4 and 8) to enrich for mitotic cells. Cells were treated with control siRNA (lanes 1-4) or siRNA targeting p300 (lanes 5-8). Western blotting demonstrated expression of the indicated proteins. B. The extracts in A were immunoprecipitated with an acetyl-lysine antibody (lanes 2-9) or an HA control antibody with HFK+HPV16 extracts (lane 1). Western blotting demonstrated the level of acetyl-lysine pull-down of the indicated proteins. C. Growing N/Tert-1+Vec and HFK+HPV16 cells were stained with the indicated antibodies and DAPI. Mitotic cells are highlighted by the white arrows.

Finally, we confirmed that E2 and TopBP1 co-localize on mitotic chromatin in HFK+HPV16 cells (Figure 6C).

## Discussion

This report describes a novel E2-TopBP1 centered mitotic complex (ECMC) that controls mitotic acetylation and expression of proteins critical for the viral life cycle, including E2 and TopBP1. The increased acetylation is due to a reduction in SIRT1 function during mitosis in an E2-TopBP1 dependent manner. In HFK+HPV16 cells there is a reduction of SIRT1 levels during mitosis, while in N/Tert-1 cells SIRT1 function is turned off as demonstrated by increased acetylation of p53 on lysine 382, a known SIRT1 target residue. SIRT1 turn off is dependent upon E2 interaction with TopBP1 as an E2 mutant that fails to bind TopBP1 (where serine 23 is changed to an alanine, S23A (22)) is unable to alter p53, TopBP1 or E2 acetylation during mitosis in N/Tert-1 or U2OS cells. During mitosis SIRT1 is unable to complex with E2 or p53 (which it can outside of mitosis) suggesting that structural changes in the E2-TopBP1 complex during mitosis control SIRT1 partner protein interactions. TopBP1 remains complexed with SIRT1 throughout mitosis, and SIRT1 can regulate the function of TopBP1 in metabolic and DNA damage pathways (26, 51). Increased acetylation of TopBP1 occurs following activation of the DNA damage response (DDR) which inactivates SIRT1 function (26); this also increases p53 acetylation, stability and function to mediate the DDR (52). The DDR is active in HPV positive cells and is critical for the viral life cycle, as is the expression of SIRT1 (25, 53–56). Other SIRT1 substrates, including NBS1 and WRN are critical for HPV life cycles (55, 57–59). During mitosis, SIRT1 condenses chromosomes due to deacetylation of histones (31), and the results presented here suggest that in HPV transformed cells there may be alternative mechanisms for carrying out this process as SIRT1 is, at least partially, inactivated. To our knowledge, we report for the first time that SIRT1 is acetylated and in E2 expressing cells this acetylation is reduced during mitosis. How the E2-TopBP1 interaction is regulating SIRT1 mitotic acetylation and function remains to be determined, but the altered SIRT1 acetylation likely contributes to the alteration in mitotic SIRT1 interacting partners and function. It is possible that mitotic kinases target the E2-TopBP1 complex to change the structure allowing for the sequestration of SIRT1 from binding partners and the blocking of its acetylation. The acetylation of SIRT1, unlike that of E2 and TopBP1, is independent of p300 and the SIRT1 acetylase could be targeted by mitotic kinases to block SIRT1 acetylation and therefore function. Future studies will focus on identifying the E2-TopBP1 dependent mechanism that regulates SIRT1 function during mitosis.

The results present a mechanism for regulating E2 stability during mitosis (Figure 7). As cells enter mitosis the ability of SIRT1 to bind and deacetylate E2 is abrogated resulting in increased K111 acetylation by p300. This promotes the recruitment of Top1 that prevents the ubiquitination of K112 resulting in enhanced E2 protein stability and expression. TopBP1 is a known interactor with ubiquitin ligases that could also be in the E2-TopBP1 complex (60). This is described in the top panel of Figure 7. However, when K111 is mutated (K111R) there is a failure to interact with Top1 allowing ubiquitination of K112 and proteasomal degradation (Figure 7, bottom panel). Undoubtedly other protein factors are involved in the E2-TopBP1 complex and future studies will seek to characterize the E2-TopBP1 complex in and out of mitosis.

**Figure 7.**
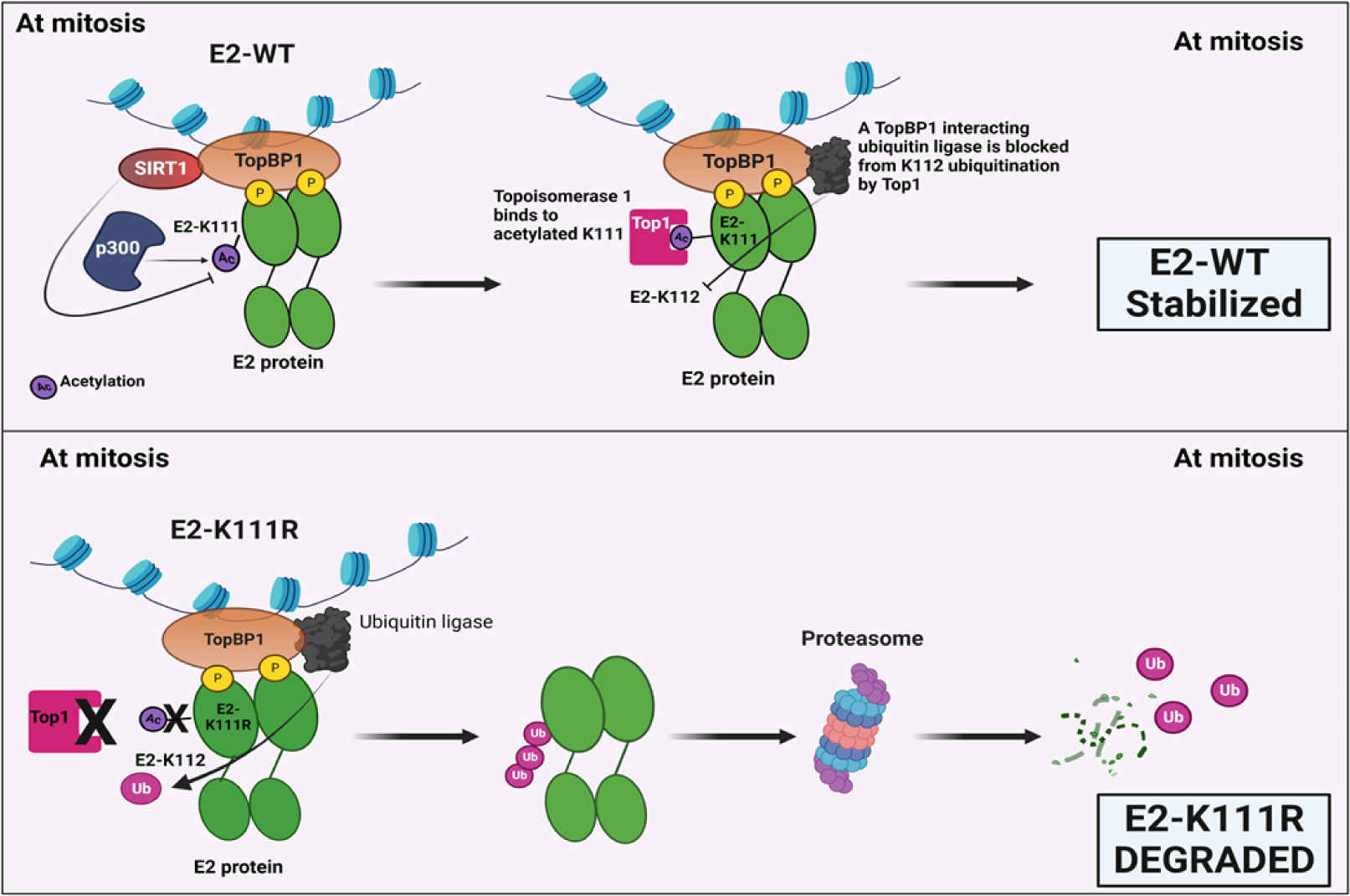
A model explaining the regulation of E2 stability during mitosis. See text for details.

We recently demonstrated that the E2-TopBP1 interaction is critical for the plasmid segregation function of E2 (21–23). This function of E2 promotes the nuclear localization of viral genomes following mitosis by allowing the viral genomes to “hitch-hike” onto the host mitotic chromatin using the E2-TopBP1 interaction. Disruption of the E2-TopBP1 interaction also resulted in a loss of E2 protein expression in HFK+HPV16 cells, promoting viral genome integration (20). In this report, we demonstrate that mutation of E2 K111 or K112 prevented the viral genome from immortalizing human foreskin keratinocytes (HFK, Table 1). This failure could be due to the aberrant expression and mitotic interaction of the E2 mutants demonstrated in this report, resulting in a loss of the viral DNA from daughter nuclei following mitosis, reducing viral nuclear DNA to a level that does not support immortalization. It is interesting that E2-K111Q can complex with mitotic chromatin and remove TopBP1. What could be occurring is that TopBP1 recruits E2-K111Q which then permanently binds to another protein (BRD4 is an obvious candidate) changing TopBP1 structure (or BRD4 structure) to remove TopBP1 from the mitotic chromatin.

Increased p53 acetylation due to inactivation of SIRT1 following activation of the DDR ordinarily promotes the ability of p53 to activate genes that arrest the cell cycle and promote DNA repair (52). Therefore, the p53 acetylation during mitosis in HFK+HPV16 cells is surprising as these cells are proliferating and must pass through mitosis to allow continued proliferation. The role of p53 in the HPV16 life cycle is not clear. E6 expression certainly degrades p53 but in cells immortalized by the full genome E6 can be spliced to E6* which removes the ability to degrade p53 (28). Recently we demonstrated robust p53 protein expression in HFK+HPV16, in HPV16 positive head and neck cancer cell lines, and in HPV16 positive head and neck cancer patient derived xenografts (29). Overexpression of full length E6 in HFK+HPV16 cells abrogated their growth (an E6 mutant unable to degrade p53 did not) suggesting that p53 may be important for the HPV16 life cycle. We have been unable to use CRISPR/Cas9 to remove p53 expression from HFK+HPV16 cells, even with target RNAs that do reduce p53 expression in non-HPV16 cells (not shown). Future studies will further investigate a possible role for p53 in the viral life cycle. p53 can directly interact with E2 and can regulate E2 replication function, therefore it is possible that p53 plays a role in the regulation of viral replication (61, 62).

The enhanced acetylation of E2 and TopBP1 during mitosis is predominantly carried out by p300. p300 is known to regulate E2 function and also to regulate E6 function (33, 44, 46, 47, 63–67) and therefore this mitotic acetylation by p300 could change the function of E2 and TopBP1.

Overall, the results identify a novel mitotic complex that is regulated by E2 interaction with TopBP1, which inhibits SIRT1 mitotic function, promoting acetylation of viral and host proteins. We have already demonstrated that the E2-TopBP1 interaction is critical for the HPV16 life cycle, and the results here also demonstrate that regulation of E2 K111 acetylation and K112 ubiquitination are also critical for HPV16 immortalization. Future studies will focus on gaining a greater understanding of this E2-TopBP1 centered mitotic complex and what the critical aspects are for the HPV16 life cycle.

## Materials and methods

### Immortalization of human foreskin keratinocytes (HFK)

Human foreskin keratinocytes (HFKs) were immortalized with HPV16 as described previously (22). Briefly, 2×10^5^ low passage (p2-p5) primary cells were plated onto collagen-coated 6cm dishes and, once at 60% confluency, were transfected with recircularized HPV16 wild-type or mutant genomes alongside neomycin resistance plasmid, pCDNA3.1. These then underwent selection, consisting of 8 days of alternate treatment with media containing G418 and 10uM Y-27632, and with mitomycin-c inactivated J2 fibroblasts. Once selected, cells were cultured in kSFM without additional drugs with mitomycin-c inactivated J2 fibroblasts, and transferred to 10 cm dishes. Once colonies were visible in the wild-type cells, media was removed and cells were washed twice with PBS. 5 mL of 0.5% crystal violet was added per dish, and then incubated shaking at room temperature for 30 mins. Dishes were washed five times with PBS and plates left to dry. Crystal violet images were scanned using the Odyssey^®^ CLx Imaging System and quantified by eye. Immortalization was done in duplicate in two donor backgrounds, individually.

### Cell culture and plasmids

Using low passage N/Tert-1 cells and U2OS cells, stably expressing HPV16 wild-type E2 (E2-WT), mutants E2-S23A (E2 with serine 23 mutated to alanine, abrogating interaction with TopBP1), E2-K111R (E2 with lysine 111 mutated to arginine), E2-K112R (E2 with lysine 112 mutated to arginine), E2-K111Q (E2 with lysine 111 mutated to glutamine), E2-K111R + K112R (E2 with lysine 111 and lysine K112 mutated to arginine) and an empty vector plasmid control, pCDNA (Vec) were generated as previously described (21, 22). N/Tert-1 cells were passaged in keratinocyte serum-free medium (K-SFM) (Invitrogen; catalog no. 37010022) supplemented with bovine pituitary extract, EGF (Invitrogen), 0.3 mM calcium chloride (Sigma; 21115) and 150 μg/ml G418 (Thermo Fisher Scientific) cultured at 37 °C in a 5% CO_2_/ 95% air atmosphere. U2OS cells were cultured in Dulbeccos Modified Eagle Medium (DMEM) (Invitrogen; catalog no. 11995065) supplemented with 10% fetal bovine serum (FBS) (R&D Systems) and 1.5 mg/mL G418 sulfate as described previously (21, 22).

Human foreskin keratinocytes (HFKs) were immortalized with HPV16 (WT) or immortalized with E6/E7 were cultured in Dermalife-K complete medium (Lifeline Cell Technology) as previously described (22). Mitomycin C-treated 3T3-J2 fibroblasts feeders were plated 24 hours prior to plating N/Tert-1-Vec or HFK cells on top of the feeders, in their respective cell culture medium and allowed to grow to 70% confluency. C33a cells were obtained from ATCC (ATCC HTB-31) and grown in DMEM Medium supplemented with 10% FBS. In all cases, cell identity was confirmed via “fingerprinting”, and cell cultures were routinely tested for mycoplasma.

All the HPV16 plasmids used in these studies have been previously described by our laboratory: HPV16 pOriLacZ (pOri16LacZ), HPV16 E1-(hemagglutinin, HA) (E1), HPV16 E2, pGL3 Basic, pGL3 Control, ptk6E2 (22, 68, 69) E2-K mutant plasmids were generated by GenScript.

### Cell synchronization

N/Tert-1 vector control cells (in K-SFM medium) and HFK cells immortalized with HPV16 genomes (HFK+HPV16) as well as HFK + E6/E7 cells (in Dermalife-K complete medium) were cultured with Mitomycin C-treated 3T3-J2 fibroblasts. Cells were plated at 5 × 10^5^ density onto 100-mm plates. The cells were treated with 2 mM thymidine diluted in their respective medium for 16 h. Cells were then washed 2 times with phosphate-buffered saline (PBS) and recovered in their respective medium. After 8 h, to block the cells at G1/S phase, a second dose of 2 mM thymidine was added, and the cells were incubated for 17 h. The cells were then washed twice with PBS and recovered as before at the following time points. For HFK cells were harvested at 0 h (G1/S phase) and 19 h (M1 phase). The above procedure was repeated in N/Tert-1 cells and U2OS cells expressing stable E2-WT, E2-S23A, E2-K111R, E2-K112R, E2-K111Q, E2-K111R+ K112R, along with plasmid control cells which were plated at a density of 5 × 10^5^ in K-SFM medium (for N/Tert-1 cells) or 3 × 10^5^ in DMEM medium (for U2OS cells) on 100-mm plates and the double thymidine blocked N/Tert-1 cells were harvested at 0 h (G1/S phase) and 16 h (M1 phase). Double thymidine blocked U2OS cells harvested at 0 h (G1/S phase) and 8 h (M1 phase). Using the harvested cells at the time points mentioned, cell lysates were prepared, and immunoblotting was carried out. Cyclin B1 antibody was used to confirm the mitosis phase in these cells using immunoblotting as described below.

### Protein isolation and immunoblotting

Cells were trypsinized, washed with 1X PBS and resuspended in 2x pellet volume protein lysis buffer (0.5% Nonidet P-40, 50 mM Tris [pH 7.8], 150 mM NaCl) supplemented with protease inhibitor (Roche Molecular Biochemicals) and phosphatase inhibitor cocktail (Sigma). Cell pellet-buffer suspension was incubated on ice for 20 min and afterwards centrifuged for 15 min at 14,000 rcf at 4 °C. Protein concentration was determined using the Bio-Rad protein estimation assay according to manufacturer’s instructions. 100 μg protein was mixed with 4x Laemmli sample buffer (Bio-Rad) and heated at 95 °C for 5 min. Protein samples were separated on Novex 4–12% Tris-glycine gel (Invitrogen) and transferred onto a nitrocellulose membrane (Bio-Rad) at 30V overnight using the wet-blot transfer method. Membranes were then blocked with Odyssey (PBS) blocking buffer (diluted 1:1 with 1X PBS) at room temperature for 1h and probed with indicated primary antibody diluted in Odyssey blocking buffer, overnight. Membranes were washed twice with PBS-Tween and an additional wash with 1X PBS and probed with the Odyssey secondary antibody (goat anti-mouse IRdye 800CW or goat anti-rabbit IRdye 680CW) (Licor) diluted in Odyssey blocking buffer at 1:10,000. Membranes were then washed as before. Membranes were imaged using the Odyssey^®^ CLx Imaging System and ImageJ was used for quantification, utilizing GAPDH as internal loading control. The following primary antibodies were used for immunoblotting in this study: monoclonal anti-E2 (B9) 1:500 (70), anti-TopBP1 1:1,000 (Bethyl, catalog no. A300-111A), anti-SIRT1 antibody 1:1,000 (Sigma, catalog no. 07-131), anti-p300 1:2,000 (Bethyl, catalog no. A300-358A), anti-Top1 1:200 (Santa Cruz; catalog no. sc-32736), anti-p53 1:1,000 (Thermo Fisher, catalog no. PA5-27822), anti-Acetyl-p53 (K382) 1:1,000 (Cell Signaling Technology, catalog no. 2525), anti-Cyclin B1 (D5C10) XP 1:1,000 (Cell Signaling Technology, catalog no. 4138) and anti-Glyceraldehyde-3-phosphate dehydrogenase (GAPDH) 1:10,000 (Santa Cruz; catalog no. sc-47724).

### Immunoprecipitation

250 μg of protein lysate from indicated cells (prepared as described above) was incubated with primary antibody of interest or a HA tag antibody (used as a negative control). 1 μg of antibody per 100 μg protein lysate was used per reaction. The protein lysate-antibody mixture was made up to a total volume of 500 μl with lysis buffer supplemented with protease inhibitors and phosphatase inhibitor cocktail and rotated end-to-end at 4°C overnight. Next day, 40 μl of prewashed protein A beads per sample (Sigma; prewashed in lysis buffer as mentioned in the manufacturer’s protocol) was added to the lysate-antibody mixture and rotated for an additional 4 h at 4°C. The samples were washed gently with 500 μl lysis buffer and centrifuged at 1,000 rcf for 3 min. This wash step was repeated three times. The bead pellet was resuspended in 4X Laemmli sample buffer (Bio-Rad), heat denatured, and centrifuged at 1,000 rcf for 3 min. Proteins were separated using a sodium dodecyl sulfate-polyacrylamide gel electrophoresis (SDS-PAGE) system and transferred onto a nitrocellulose membrane before probing for the presence of E2, TopBP1, p300, SIRT1, Top1, p53 as per the western blotting protocol described above.

### RNA isolation and SYBR green reverse transcription

RNA was isolated using the SV Total RNA isolation system (Promega) following the manufacturer’s instructions. We reverse transcribed 2 μg of RNA into cDNA using the high-capacity reverse transcription kit (Applied Biosystems). cDNA was then processed for qPCR.

### Real-time PCR (qPCR)

qPCR was performed on cDNA isolated, as described above. DNA and relevant primers were mixed with PowerUp SYBR green master mix (Applied Biosystems), and using SYBR green reagent, real-time PCR was performed in the 7500 Fast real-time PCR system. Expression was quantified as relative quantity over GAPDH using the 2−ΔΔCT method. Primers used are as follows. HPV16 E2 F, 5′-ATGGAGACTCTTTGCCAACG-3′; HPV16 E2 R, 5′-TCATATAGACATAAATCCAG-3′; TopBP1 F, 5′-TGAGTGTGCCAAGAGATGGAA-3′; TopBP1 R, 5′-TGCTTCTGGTCTAGGTTCTGT-3′; SIRT1 F, 5′-CAGTGTCATGGTTCCTTTGC-3′; SIRT1 R, 5′-CACCGAGGAACTACCTGAT-3′; p53 F, 5′-GAGGTTGGCTCTGACTGTACC-3’; p53 R, 5’-TCCGTCCCAGTAGATTACCAC-3’; Glyceraldehyde-3-phosphate dehydrogenase (GAPDH) F, 5′-GGAGCGAGATCCCTCCAAAAT-3′ and GAPDH R, 5′-GGCTGTTGTCATACTTCTCATGG-3′.

### MG132 proteasomal inhibitor treatment

N/Tert-1 cells expressing stable E2-WT or E2-K111R, E2-1112R, E2-111Q, E2-K111R+K1112R mutants were plated in KSFM media. Next, double thymidine treatment was carried out as described above. At 15h post second thymidine recovery, cells were treated with 10μM of MG132 (Z-Leu-Leu-Leu-al; Sigma, catalog no. C2211). Cells were then harvested at 1h after MG132 treatment (corresponds to 16h post second thymidine recovery; peak M1 phase in N/Tert-1 cells) and at 2h after MG132 treatment (corresponds to 18h post second thymidine recovery). Immunoblotting was carried out to detect E2 and TopBP1 levels.

### Ubiquitin trap

Protein lysates from the above-mentioned cell synchronized and MGM132 treated N/Tert-1 cells were harvested using NP40 protein lysis buffer. 300 μg of protein lysate was incubated with 25 μl ChromoTek Ubiquitin-Trap Agarose slurry (Proteintech) equilibrated in NP40 protein lysis buffer and placed at 4°C for 1 h with continual end-to-end rotation. The protein-bound Ubiquitin-Trap Agarose were washed three times in the NP40 lysis buffer by centrifugation at 1,000 rcf for 3 min and resuspended in 4× Laemmli sample buffer (Bio-Rad), heat denatured, and centrifuged at 1,000 rcf for 3 min. The supernatant was gel electrophoresed using an SDS-PAGE system which was later transferred onto a nitrocellulose membrane using wet-blot transfer method. The membrane was probed with an E2 antibody to detect ubiquitinated E2.

### Immunofluorescence

U2OS cells expressing stable E2-WT, E2-K111R, E2-1112R, E2-111Q, E2-K111R+K1112R and pcDNA empty vector plasmid control were plated on acid-washed, poly-l-lysine-coated coverslips, in a six-well plate at a density of 2 × 10^5^ cells/well with 5 ml DMEM with 10% FBS. After 48 h, the cells were washed twice with PBS, fixed, and stained as previously described (21, 22). The primary antibodies used are as follows: HPV16 E2 B9 monoclonal antibody 1:500; TopBP1, 1:1,000 (Bethyl Laboratories, catalog no. A300-111A). The cells were washed and incubated with secondary antibodies Alexa Fluor 488 goat anti-mouse (Thermo Fisher, catalog no. A-11001) and Alexa Fluor 594 goat anti-rabbit (Thermo Fisher, catalog no. A-11037) diluted at 1:1,000. The wash step was repeated. Nuclear DNA was stained with 4’,6-diamidino-2-phenylindole (DAPI) (Santa Cruz, catalog no. sc-3598) and the coverslips were mounted on a glass slide using Vectashield mounting medium (ThermoFisher). Images were visualized, captured using a Zeiss LSM700 laser scanning confocal microscope and analyzed and quantitated using Zen LE software and Keyence analyzing system (BZ-X810).

### Small interfering RNA (siRNA) treatment

HFK cells or N/Tert-1 cells were plated on top of Mitomycin C-treated 3T3-J2 fibroblasts feeders in 100-mm plates in their respective media. The next day, cells were transfected with 10 µM of the siRNA mentioned below. 10 µM of a “non-targeting” control, MISSION^®^ siRNA Universal Negative Control (Sigma-Aldrich; catalog no. SIC001), was used in our experiments. siRNA knockdown was carried out following the protocol from Lipofectamine™ RNAiMAX transfection (Invitrogen, catalog no. 13778-100). Cells were harvested 48 hours post transfection and using the protocol as described above, immunoblotting for the protein of interest was done to confirm the knockdown. All siRNAs were purchased from Sigma-Aldrich: siRNA p300-A,5’-TTGGACTACCCTATCAAGTAA-3’; siRNA p300-B,5’-GACUACCCUAUCAAGUAA-3’.

### DNA Replication Assay

On 100-mm dishes, C33a cells were plated at 5×10^5^ in DMEM + 10% FBS. The following day, plasmid DNA was transfected using the calcium phosphate method (71). Three days post-transfection, low molecular weight DNA was extracted using the Hirt method as previously described (72). The sample was extracted twice with phenol:chloroform:isoamyl alcohol (25:24:1) and precipitated with ethanol. Following centrifugation, the DNA pellet was washed with 70% ethanol, dried and resuspended in a total of 150 µL water. Forty-two microliters of sample were digested with DpnI (New England Biolabs) overnight to remove unreplicated pOri16LacZ; the sample was then digested with ExoIII (New England Biolabs) for 1 hr. Replication was determined by real-time PCR, as described previously (69).

### Transcription assay

C33a cells were plated at 5 × 10^5^ on 100-mm plates in DMEM medium with 10% FBS. Next day, the cells were transfected with either 1 μg HPV16 E2 plasmids (WT or E2-K mutants) and 1 μg ptk6E2-luc or 1 μg ptk6E2-luc alone using calcium phosphate transfection method. Briefly, the cells were harvested 72 h post transfection utilizing the Promega reporter lysis buffer and analyzed for luciferase using the Promega luciferase assay system (catalog no. E1500). Concentrations were normalized to protein levels, as measured by the Bio-Rad protein estimation assay mentioned above. Relative fluorescence units were measured using the BioTek Synergy H1 hybrid reader.

### Statistical analysis

All the experiments were carried out in triplicates in each of the mentioned cell lines and quantitation of the results represented as mean ± standard error (SE). Significance was determined using a student’s t-test.

## Acknowledgements

IMM’s work is supported by US NIH grant R01DE029471.

## Notes

### Competing Interest Statement

The authors have declared no competing interest.

## References

1. Doorbar J, Quint W, Banks L, Bravo IG, Stoler M, Broker TR, Stanley MA. 2012. The biology and life-cycle of human papillomaviruses. Vaccine 30 Suppl 5:F55–70.

2. zur Hausen H. 2009. Papillomaviruses in the causation of human cancers - a brief historical account. Virology 384:260–265.

3. Galloway DA, Laimins LA. 2015. Human papillomaviruses: shared and distinct pathways for pathogenesis. Curr Opin Virol 14:87–92.

4. Hoppe-Seyler K, Bossler F, Braun JA, Herrmann AL, Hoppe-Seyler F. 2018. The HPV E6/E7 Oncogenes: Key Factors for Viral Carcinogenesis and Therapeutic Targets. Trends in microbiology 26:158–168.

5. McBride AA. 2013. The Papillomavirus E2 proteins. Virology 445:57–79.

6. Bergvall M, Melendy T, Archambault J. 2013. The E1 proteins. Virology 445:35–56.

7. Masterson PJ, Stanley MA, Lewis AP, Romanos MA. 1998. A C-terminal helicase domain of the human papillomavirus E1 protein binds E2 and the DNA polymerase alpha-primase p68 subunit. Journal of virology 72:7407–7419.

8. Benson JD, Howley PM. 1995. Amino-terminal domains of the bovine papillomavirus type 1 E1 and E2 proteins participate in complex formation. Journal of virology 69:4364–4372.

9. Bouvard V, Storey A, Pim D, Banks L. 1994. Characterization of the human papillomavirus E2 protein: evidence of trans-activation and trans-repression in cervical keratinocytes. The EMBO journal 13:5451–5459.

10. Evans MR, James CD, Bristol ML, Nulton TJ, Wang X, Kaur N, White EA, Windle B, Morgan IM. 2019. Human Papillomavirus 16 E2 Regulates Keratinocyte Gene Expression Relevant to Cancer and the Viral Life Cycle. J Virol 93.

11. Schweiger MR, Ottinger M, You J, Howley PM. 2007. Brd4-independent transcriptional repression function of the papillomavirus e2 proteins. Journal of virology 81:9612–9622.

12. Schweiger MR, You J, Howley PM. 2006. Bromodomain protein 4 mediates the papillomavirus E2 transcriptional activation function. Journal of virology 80:4276–4285.

13. McBride AA, Sakakibara N, Stepp WH, Jang MK. 2012. Hitchhiking on host chromatin: how papillomaviruses persist. Biochimica et biophysica acta 1819:820–825.

14. You J, Schweiger MR, Howley PM. 2005. Inhibition of E2 binding to Brd4 enhances viral genome loss and phenotypic reversion of bovine papillomavirus-transformed cells. Journal of virology 79:14956–14961.

15. You J, Croyle JL, Nishimura A, Ozato K, Howley PM. 2004. Interaction of the bovine papillomavirus E2 protein with Brd4 tethers the viral DNA to host mitotic chromosomes. Cell 117:349–360.

16. Adam S, Rossi SE, Moatti N, De Marco Zompit M, Xue Y, Ng TF, Álvarez-Quilón A, Desjardins J, Bhaskaran V, Martino G, Setiaputra D, Noordermeer SM, Ohsumi TK, Hustedt N, Szilard RK, Chaudhary N, Munro M, Veloso A, Melo H, Yin SY, Papp R, Young JTF, Zinda M, Stucki M, Durocher D. 2021. The CIP2A-TOPBP1 axis safeguards chromosome stability and is a synthetic lethal target for BRCA-mutated cancer. Nat Cancer 2:1357–1371.

17. Wang X, Helfer CM, Pancholi N, Bradner JE, You J. 2013. Recruitment of Brd4 to the human papillomavirus type 16 DNA replication complex is essential for replication of viral DNA. Journal of virology 87:3871–84.

18. Helfer CM, Wang R, You J. 2013. Analysis of the papillomavirus E2 and bromodomain protein Brd4 interaction using bimolecular fluorescence complementation. PLoS One 8:e77994.

19. McPhillips MG, Oliveira JG, Spindler JE, Mitra R, McBride aa. 2006. Brd4 is required for e2-mediated transcriptional activation but not genome partitioning of all papillomaviruses. Journal of virology 80:9530–43.

20. Prabhakar AT, James CD, Fontan CT, Otoa R, Wang X, Bristol ML, Hill RD, Dubey A, Morgan IM. 2023. Human Papillomavirus 16 E2 Interaction with TopBP1 Is Required for E2 and Viral Genome Stability during the Viral Life Cycle. J Virol 97:e0006323.

21. Prabhakar AT, James CD, Das D, Fontan CT, Otoa R, Wang X, Bristol ML, Morgan IM. 2022. Interaction with TopBP1 Is Required for Human Papillomavirus 16 E2 Plasmid Segregation/Retention Function during Mitosis. J Virol 96:e0083022.

22. Prabhakar AT, James CD, Das D, Otoa R, Day M, Burgner J, Fontan CT, Wang X, Glass SH, Wieland A, Donaldson MM, Bristol ML, Li R, Oliver AW, Pearl LH, Smith BO, Morgan IM. 2021. CK2 Phosphorylation of Human Papillomavirus 16 E2 on Serine 23 Promotes Interaction with TopBP1 and Is Critical for E2 Interaction with Mitotic Chromatin and the Viral Life Cycle. mBio doi:10.1128/mBio.01163-21:e0116321.

23. Prabhakar AT, James CD, Fontan CT, Otoa R, Wang X, Bristol ML, Yeager C, Hill RD, Dubey A, Wu SY, Chiang CM, Morgan IM. 2023. Direct interaction with the BRD4 carboxyl-terminal motif (CTM) and TopBP1 is required for human papillomavirus 16 E2 association with mitotic chromatin and plasmid segregation function. J Virol doi:10.1128/jvi.00782-23:e0078223.

24. Das D, Smith N, Wang X, Morgan IM. 2017. The Deacetylase SIRT1 Regulates the Replication Properties of Human Papillomavirus 16 E1 and E2. Journal of virology 91:e00102–e00117.

25. Langsfeld ES, Bodily JM, Laimins LA. 2015. The Deacetylase Sirtuin 1 Regulates Human Papillomavirus Replication by Modulating Histone Acetylation and Recruitment of DNA Damage Factors NBS1 and Rad51 to Viral Genomes. PLoS pathogens 11:e1005181.

26. Liu T, Lin YH, Leng W, Jung SY, Zhang H, Deng M, Evans D, Li Y, Luo K, Qin B, Qin J, Yuan J, Lou Z. 2014. A divergent role of the SIRT1-TopBP1 axis in regulating metabolic checkpoint and DNA damage checkpoint. Molecular cell 56:681–695.

27. Wang R-H, Lahusen TJ, Chen Q, Xu X, Jenkins LMM, Leo E, Fu H, Aladjem M, Pommier Y, Appella E, Deng C-X. 2014. SIRT1 deacetylates TopBP1 and modulates intra-S-phase checkpoint and DNA replication origin firing. International journal of biological sciences 10:1193–202.

28. Pim D, Massimi P, Banks L. 1997. Alternatively spliced HPV-18 E6* protein inhibits E6 mediated degradation of p53 and suppresses transformed cell growth. Oncogene 15:257–64.

29. Fontan CT, James CD, Prabhakar AT, Bristol ML, Otoa R, Wang X, Karimi E, Rajagopalan P, Basu D, Morgan IM. 2022. A Critical Role for p53 during the HPV16 Life Cycle. Microbiol Spectr doi:10.1128/spectrum.00681-22:e0068122.

30. Gandhi S, Mitterhoff R, Rapoport R, Farago M, Greenberg A, Hodge L, Eden S, Benner C, Goren A, Simon I. 2022. Mitotic H3K9ac is controlled by phase-specific activity of HDAC2, HDAC3, and SIRT1. Life Sci Alliance 5.

31. Fatoba ST, Okorokov AL. 2011. Human SIRT1 associates with mitotic chromatin and contributes to chromosomal condensation. Cell Cycle 10:2317–22.

32. Thomas Y, Androphy EJ. 2018. Human Papillomavirus Replication Regulation by Acetylation of a Conserved Lysine in the E2 Protein. J Virol 92.

33. Thomas Y, Androphy EJ. 2019. Acetylation of E2 by P300 Mediates Topoisomerase Entry at the Papillomavirus Replicon. J Virol 93.

34. Trivedi P, Steele CD, Au FKC, Alexandrov LB, Cleveland DW. 2023. Mitotic tethering enables inheritance of shattered micronuclear chromosomes. Nature 618:1049–1056.

35. Lin YF, Hu Q, Mazzagatti A, Valle-Inclán JE, Maurais EG, Dahiya R, Guyer A, Sanders JT, Engel JL, Nguyen G, Bronder D, Bakhoum SF, Cortés-Ciriano I, Ly P. 2023. Mitotic clustering of pulverized chromosomes from micronuclei. Nature 618:1041–1048.

36. Gelot C, Kovacs MT, Miron S, Mylne E, Haan A, Boeffard-Dosierre L, Ghouil R, Popova T, Dingli F, Loew D, Guirouilh-Barbat J, Del Nery E, Zinn-Justin S, Ceccaldi R. 2023. Polθ is phosphorylated by PLK1 to repair double-strand breaks in mitosis. Nature 621:415–422.

37. De Marco Zompit M, Esteban MT, Mooser C, Adam S, Rossi SE, Jeanrenaud A, Leimbacher PA, Fink D, Shorrocks AK, Blackford AN, Durocher D, Stucki M. 2022. The CIP2A-TOPBP1 complex safeguards chromosomal stability during mitosis. Nat Commun 13:4143.

38. Bagge J, Oestergaard VH, Lisby M. 2020. Functions of TopBP1 in preserving genome integrity during mitosis. Semin Cell Dev Biol doi:10.1016/j.semcdb.2020.08.009.

39. Leimbacher PA, Jones SE, Shorrocks AK, de Marco Zompit M, Day M, Blaauwendraad J, Bundschuh D, Bonham S, Fischer R, Fink D, Kessler BM, Oliver AW, Pearl LH, Blackford AN, Stucki M. 2019. MDC1 Interacts with TOPBP1 to Maintain Chromosomal Stability during Mitosis. Mol Cell 74:571–583.e8.

40. Gallina I, Christiansen SK, Pedersen RT, Lisby M, Oestergaard VH. 2016. TopBP1-mediated DNA processing during mitosis. Cell Cycle 15:176–83.

41. Broderick R, Nieminuszczy J, Blackford AN, Winczura A, Niedzwiedz W. 2015. TOPBP1 recruits TOP2A to ultra-fine anaphase bridges to aid in their resolution. Nature communications 6:6572–6572.

42. Pedersen RT, Kruse T, Nilsson J, Oestergaard VH, Lisby M. 2015. TopBP1 is required at mitosis to reduce transmission of DNA damage to G1 daughter cells. The Journal of cell biology 210:565–582.

43. Germann SM, Schramke V, Pedersen RT, Gallina I, Eckert-Boulet N, Oestergaard VH, Lisby M. 2014. TopBP1/Dpb11 binds DNA anaphase bridges to prevent genome instability. The Journal of cell biology 204:45–59.

44. Quinlan EJ, Culleton SP, Wu SY, Chiang CM, Androphy EJ. 2013. Acetylation of conserved lysines in bovine papillomavirus E2 by p300. Journal of virology 87:1497–1507.

45. Wang X, Naidu SR, Sverdrup F, Androphy EJ. 2009. Tax1BP1 interacts with papillomavirus E2 and regulates E2-dependent transcription and stability. J Virol 83:2274–84.

46. Peng YC, Breiding DE, Sverdrup F, Richard J, Androphy EJ. 2000. AMF-1/Gps2 binds p300 and enhances its interaction with papillomavirus E2 proteins. J Virol 74:5872–9.

47. Kruppel U, Muller-Schiffmann A, Baldus SE, Smola-Hess S, Steger G. 2008. E2 and the co-activator p300 can cooperate in activation of the human papillomavirus type 16 early promoter. Virology 377:151–159.

48. Rehtanz M, Schmidt HM, Warthorst U, Steger G. 2004. Direct interaction between nucleosome assembly protein 1 and the papillomavirus E2 proteins involved in activation of transcription. Mol Cell Biol 24:2153–68.

49. Müller A, Ritzkowsky A, Steger G. 2002. Cooperative activation of human papillomavirus type 8 gene expression by the E2 protein and the cellular coactivator p300. J Virol 76:11042–53.

50. Lill NL, Grossman SR, Ginsberg D, DeCaprio J, Livingston DM. 1997. Binding and modulation of p53 by p300/CBP coactivators. Nature 387:823–7.

51. Wang RH, Lahusen TJ, Chen Q, Xu X, Jenkins LM, Leo E, Fu H, Aladjem M, Pommier Y, Appella E, Deng CX. 2014. SIRT1 deacetylates TopBP1 and modulates intra-S-phase checkpoint and DNA replication origin firing. International journal of biological sciences 10:1193–1202.

52. Vaziri H, Dessain SK, Ng Eaton E, Imai SI, Frye RA, Pandita TK, Guarente L, Weinberg RA. 2001. hSIR2(SIRT1) functions as an NAD-dependent p53 deacetylase. Cell 107:149–159.

53. Moody CA, Laimins LA. 2009. Human papillomaviruses activate the ATM DNA damage pathway for viral genome amplification upon differentiation. PLoS pathogens 5:e1000605.

54. Chappell WH, Gautam D, Ok ST, Johnson BA, Anacker DC, Moody CA. 2015. Homologous Recombination Repair Factors, Rad51 and BRCA1, are Necessary for Productive Replication of Human Papillomavirus 31. Journal of virology doi:JVI.02495-15 [pii].

55. Anacker DC, Gautam D, Gillespie KA, Chappell WH, Moody CA. 2014. Productive replication of human papillomavirus 31 requires DNA repair factor Nbs1. Journal of virology 88:8528–8544.

56. Gillespie Ka, Mehta KP, Laimins La, Moody Ca. 2012. Human papillomaviruses recruit cellular DNA repair and homologous recombination factors to viral replication centers. Journal of virology 86:9520–6.

57. Yuan Z, Zhang X, Sengupta N, Lane WS, Seto E. 2007. SIRT1 regulates the function of the Nijmegen breakage syndrome protein. Molecular cell 27:149–162.

58. Das D, Bristol ML, Smith NW, James CD, Wang X, Pichierri P, Morgan IM. 2019. Werner Helicase Control of Human Papillomavirus 16 E1-E2 DNA Replication Is Regulated by SIRT1 Deacetylation. MBio 10.

59. 59. James CD, Das D, Morgan EL, Otoa R, Macdonald A, Morgan IM. 2020. Werner Syndrome Protein (WRN) Regulates Cell Proliferation and the Human Papillomavirus 16 Life Cycle during Epithelial Differentiation. mSphere 5.

60. Honda Y, Tojo M, Matsuzaki K, Anan T, Matsumoto M, Ando M, Saya H, Nakao M. 2002. Cooperation of HECT-domain ubiquitin ligase hHYD and DNA topoisomerase II-binding protein for DNA damage response. J Biol Chem 277:3599–605.

61. Brown C, Kowalczyk AM, Taylor ER, Morgan IM, Gaston K. 2008. P53 represses human papillomavirus type 16 DNA replication via the viral E2 protein. Virology journal 5:5–422X-5-5.

62. Parish JL, Kowalczyk A, Chen H-t, Roeder GE, Sessions R, Buckle M, Gaston K. 2006. E2 Proteins from High- and Low-Risk Human Papillomavirus Types Differ in Their Ability To Bind p53 and Induce Apoptotic Cell Death E2 Proteins from High- and Low-Risk Human Papillomavirus Types Differ in Their Ability To Bind p53 and Induce Apoptotic Cell. doi:10.1128/JVI.80.9.4580.

63. Xie X, Piao L, Bullock BN, Smith A, Su T, Zhang M, Teknos TN, Arora PS, Pan Q. 2014. Targeting HPV16 E6-p300 interaction reactivates p53 and inhibits the tumorigenicity of HPV-positive head and neck squamous cell carcinoma. Oncogene 33:1037–1046.

64. White EA, Kramer RE, Tan MJ, Hayes SD, Harper JW, Howley PM. 2012. Comprehensive analysis of host cellular interactions with human papillomavirus E6 proteins identifies new E6 binding partners and reflects viral diversity. Journal of virology 86:13174–13186.

65. Muench P, Probst S, Schuetz J, Leiprecht N, Busch M, Wesselborg S, Stubenrauch F, Iftner T. 2010. Cutaneous papillomavirus E6 proteins must interact with p300 and block p53-mediated apoptosis for cellular immortalization and tumorigenesis. Cancer research 70:6913–6924.

66. Patel D, Huang SM, Baglia LA, McCance DJ. 1999. The E6 protein of human papillomavirus type 16 binds to and inhibits co-activation by CBP and p300. The EMBO journal 18:5061–5072.

67. Zimmermann H, Degenkolbe R, Bernard HU, O’Connor MJ. 1999. The human papillomavirus type 16 E6 oncoprotein can down-regulate p53 activity by targeting the transcriptional coactivator CBP/p300. Journal of virology 73:6209–6219.

68. Gauson EJ, Donaldson MM, Dornan ES, Wang X, Bristol M, Bodily JM, Morgan IM. 2015. Evidence supporting a role for TopBP1 and Brd4 in the initiation but not continuation of human papillomavirus 16 E1/E2-mediated DNA replication. J Virol 89:4980–91.

69. Taylor ER, Morgan IM. 2003. A novel technique with enhanced detection and quantitation of HPV-16 E1- and E2-mediated DNA replication. Virology 315:103–109.

70. Wieland A, Patel MR, Cardenas MA, Eberhardt CS, Hudson WH, Obeng RC, Griffith CC, Wang X, Chen ZG, Kissick HT, Saba NF, Ahmed R. 2020. Defining HPV-specific B cell responses in patients with head and neck cancer. Nature doi:10.1038/s41586-020-2931-3.

71. Kingston RE, Chen CA, Rose JK. 2003. Calcium Phosphate Transfection. Current Protocols in Molecular Biology 63:9.1.1–9.1.11.

72. Boner W, Taylor ER, Tsirimonaki E, Yamane K, Campo MS, Morgan IM. 2002. A Functional interaction between the human papillomavirus 16 transcription/replication factor E2 and the DNA damage response protein TopBP1. J Biol Chem 277:22297–303.

