## Supplementary Figures 1-10 for "A human papillomavirus 16 E2-TopBP1 dependent SIRT1-p300 acetylation switch regulates mitotic viral and human protein levels"

Figure S1

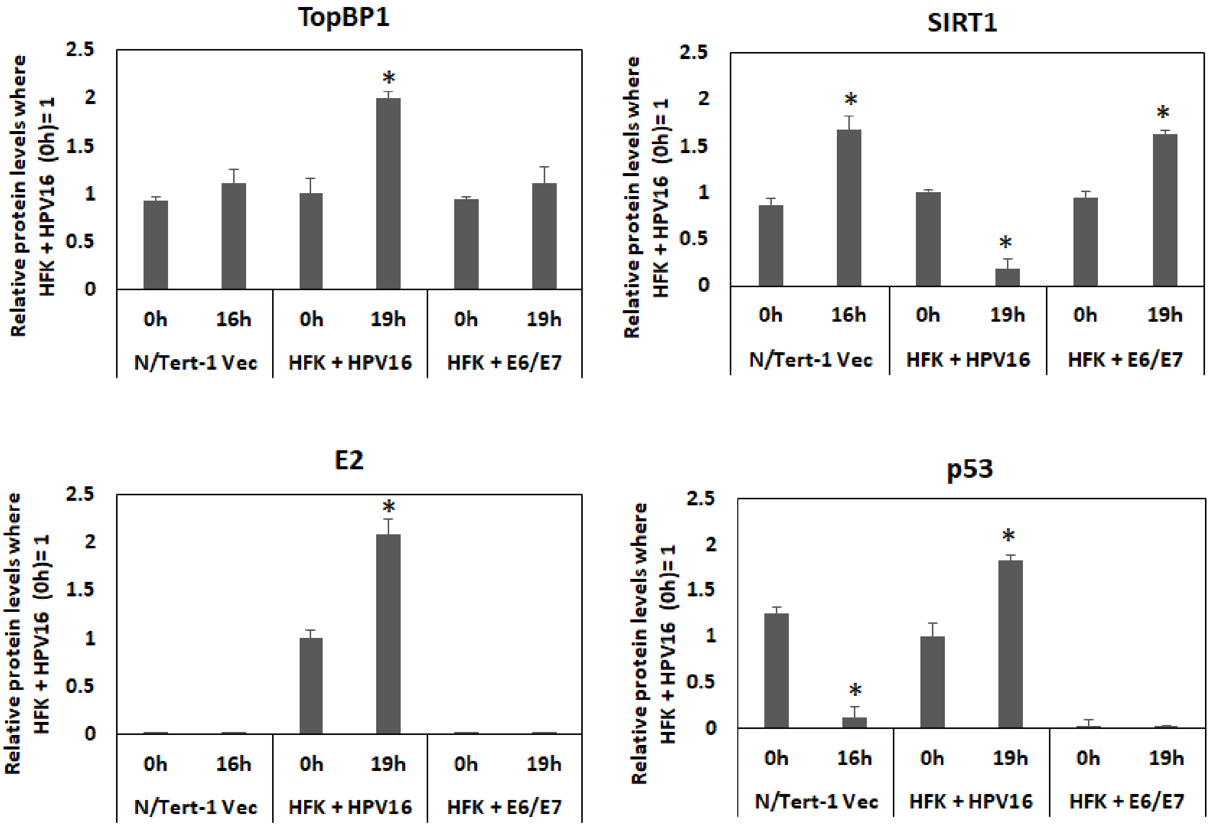


Figure S1. Repeat blots from Figure 1A were carried out and protein levels quantitated. Significant differences between 0h and 16h (N/Tert-1-Vec) and 0h and 19h (HFK+HPV16) are highlighted with *, p-value < 0.05.

Figure S2 – N/Tert-1 cells

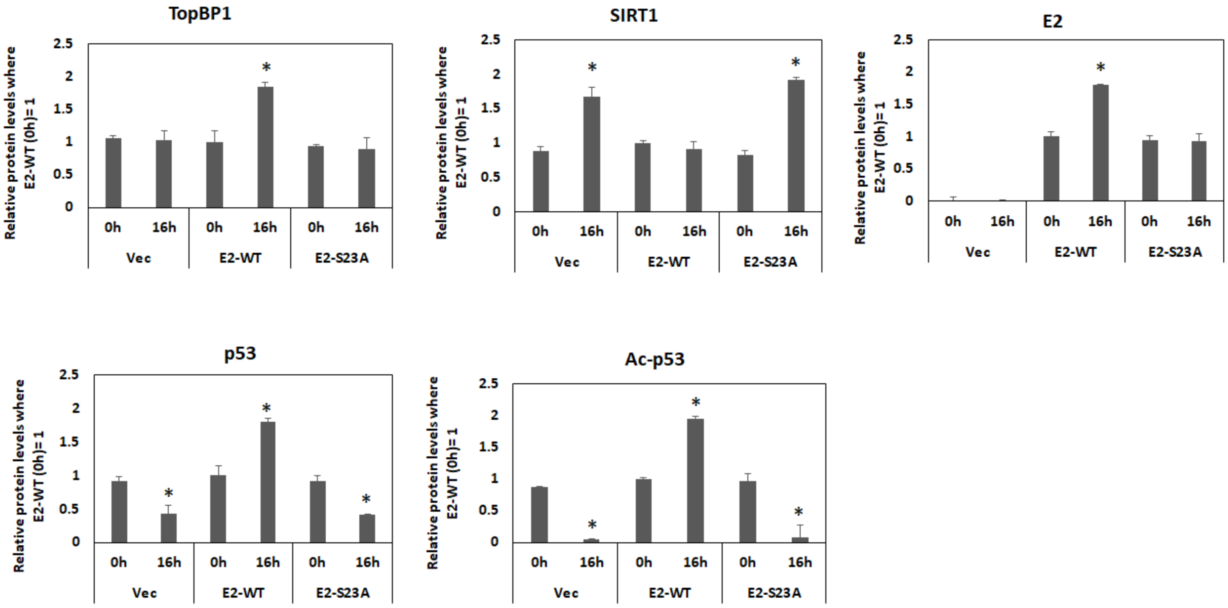


Figure S2. Repeat blots from Figure 2A were carried out and protein levels quantitated. Significant differences between 0h and 16h are highlighted with *, p-value < 0.05.

Figure S3 – U2OS cells

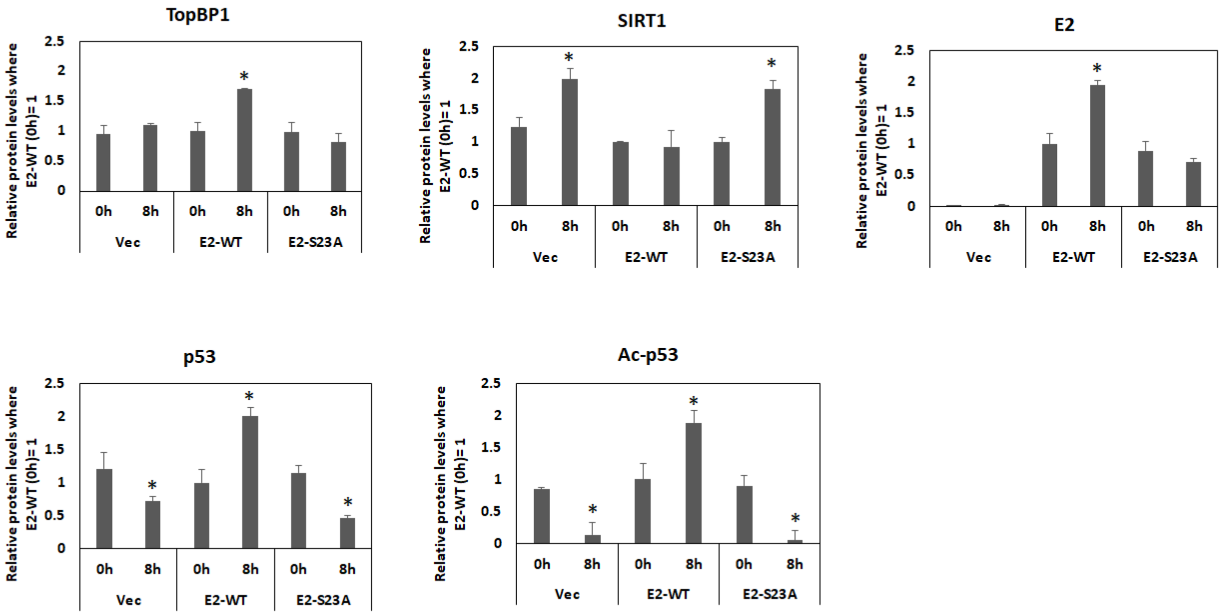


Figure S3. Repeat blots from Figure 2D were carried out and protein levels quantitated. Significant differences between 0h and 16h are highlighted with *, p-value < 0.05.

Figure S4 – N/Tert-1 cells

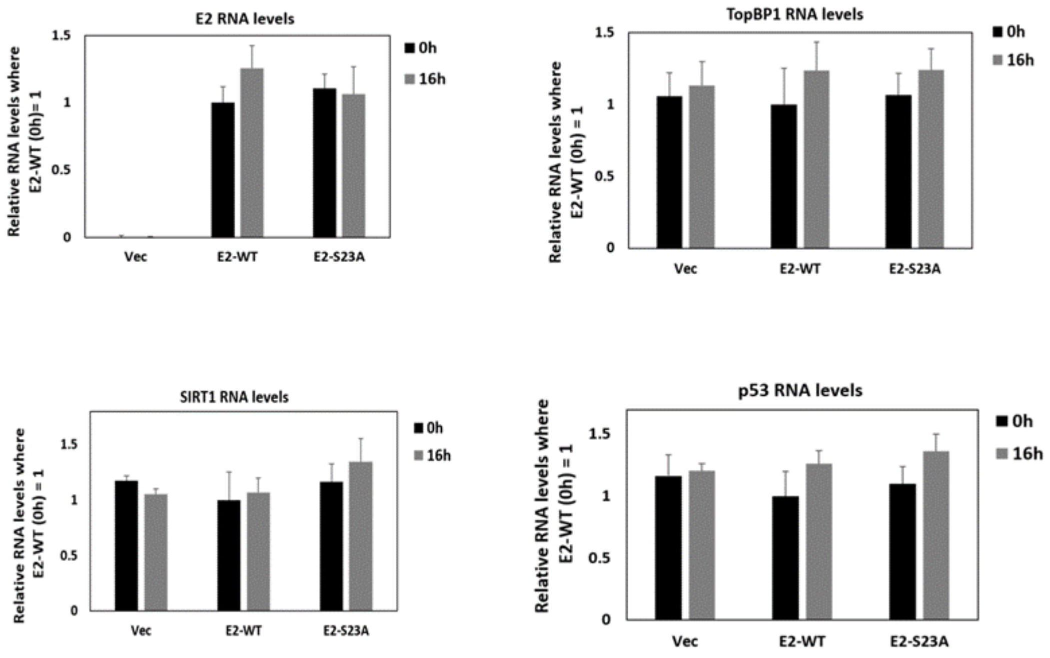


Figure S4. RNA was prepared from cells treated as in Figure 2A. The results represent a summary of three independent RNA samples and there were no significant differences.

Figure S5 – U2OS cells

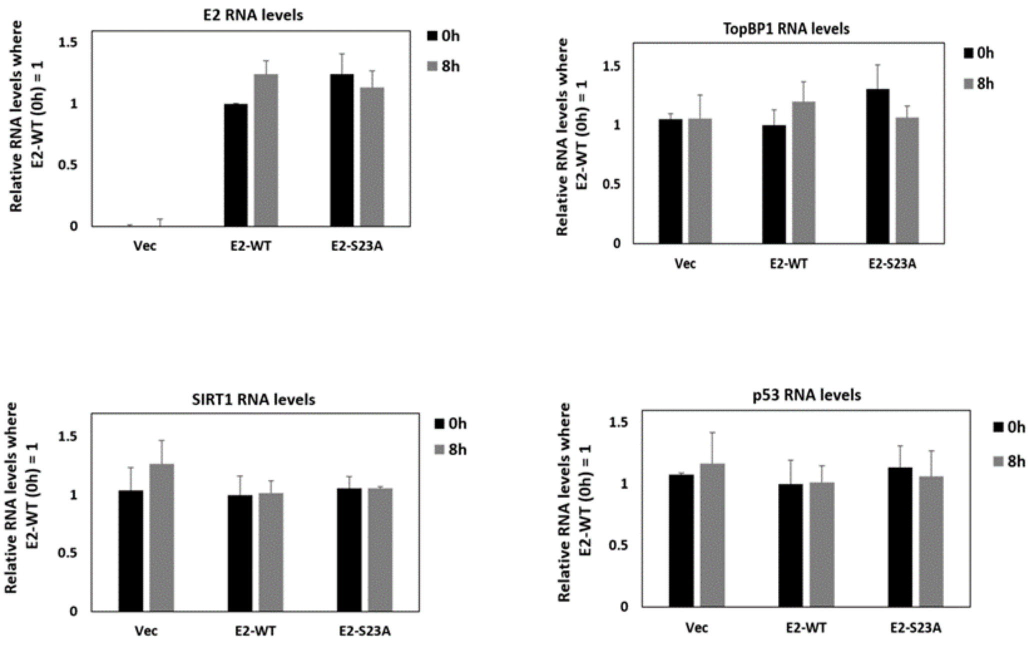


Figure S5. RNA was prepared from cells treated as in Figure 2D. The results represent a summary of three independent RNA samples and there were no significant differences.

Figure S6 – N/Tert-1 cells

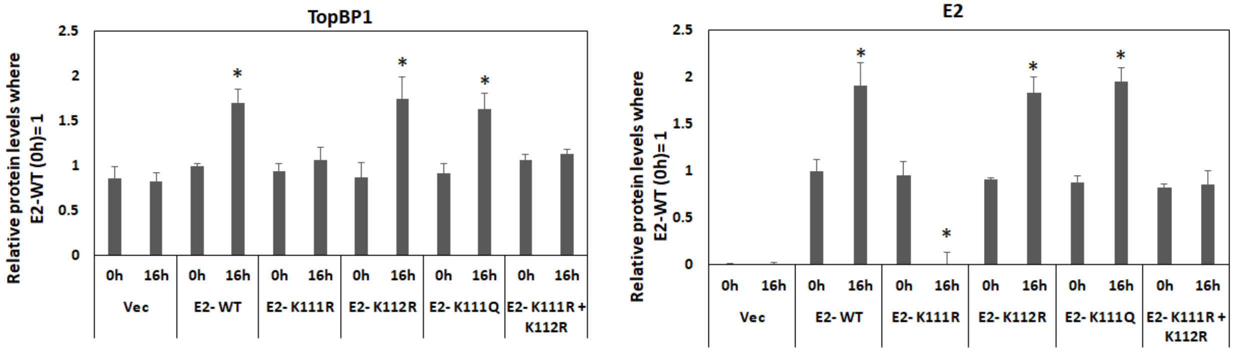


Figure S6. Repeat blots from Figure 3B were carried out and protein levels quantitated. Significant differences between 0h and 16h are highlighted with *, p-value < 0.05.

Figure S7 – N/Tert-1 cells

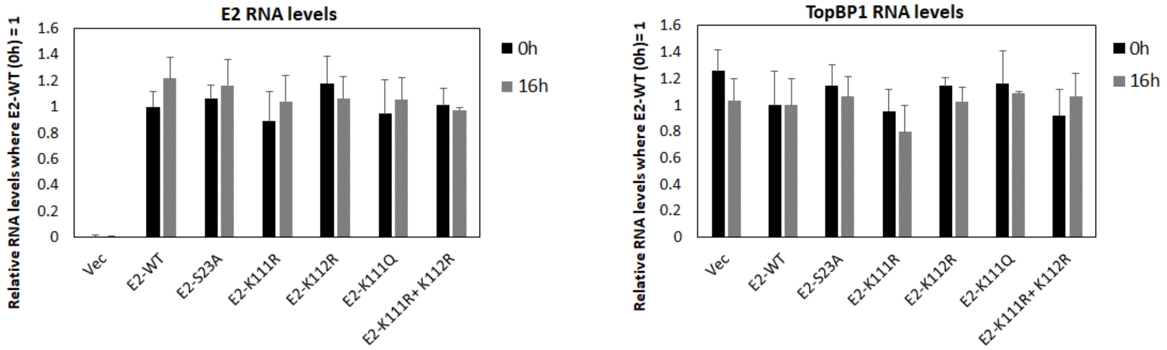


Figure S7. RNA was prepared from cells treated as in Figure 3B. The results represent a summary of three independent RNA samples and there were no significant differences.

Figure S8 – C33A cells

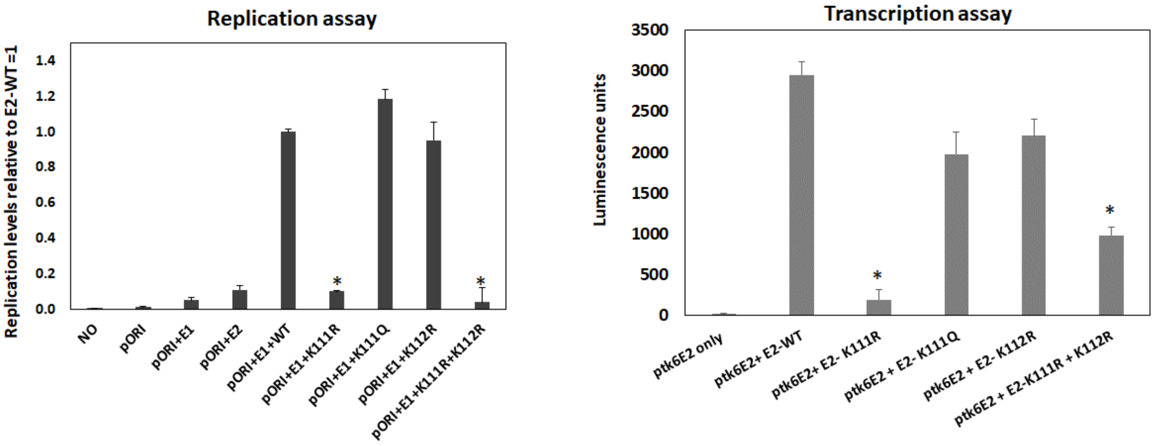


Figure S8. Transcription and replication properties of E2 proteins in C33a cells. Transcription and replication assays were carried out as described previously (1) and the results represent the summary of three independent experiments. Statistical differences from E2-WT are represented with *, p-value < 0.05.

Figure S9

A

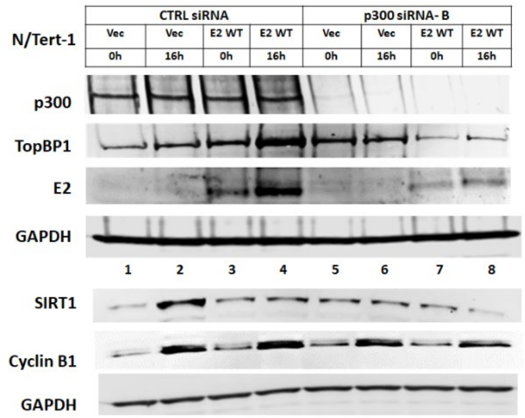


B

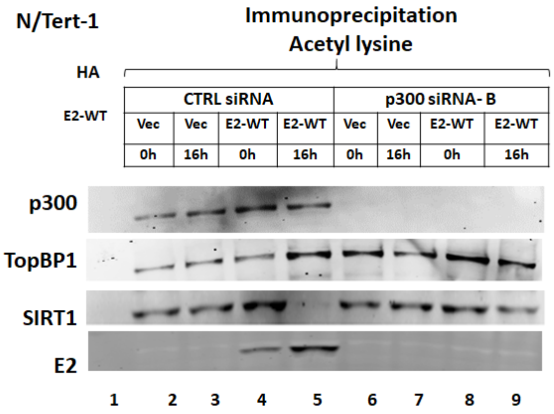


Figure S9. Repeat of Figure 5 with another p300 siRNA.

Figure S10

A B

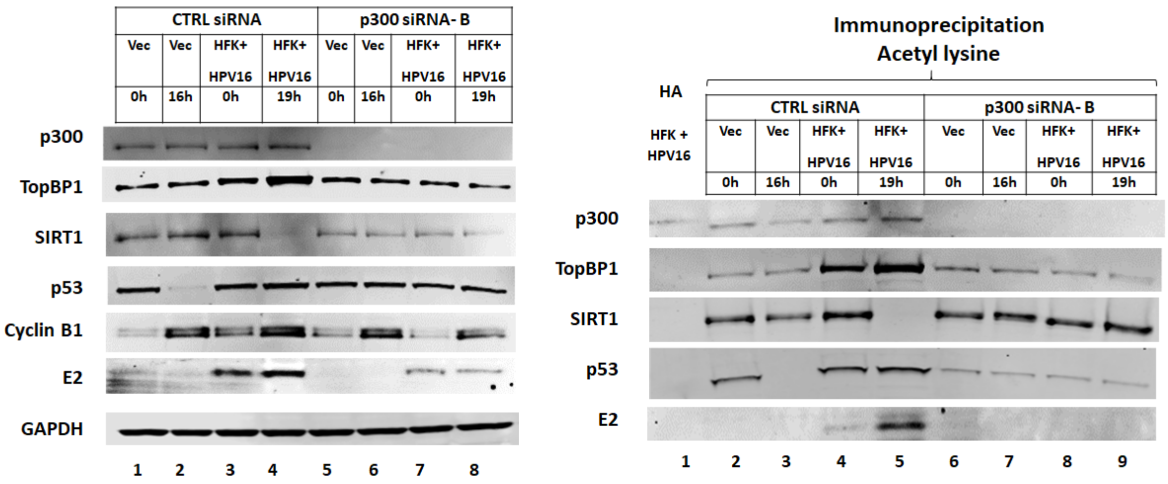


Figure S10. Repeat of Figure 6 with another p300 siRNA.


1. Gauson EJ, Donaldson MM, Dornan ES, Wang X, Bristol M, Bodily JM, Morgan IM. 2015. Evidence supporting a role for TopBP1 and Brd4 in the initiation but not continuation of human papillomavirus 16 E1/E2 mediated DNA replication. Journal of virology 89:17684-17699.
